# Single-cell transcriptomics identifies altered neutrophil dynamics and accentuated T-cell cytotoxicity in tobacco flavored e-cigarette exposed mouse lungs

**DOI:** 10.1101/2025.02.17.638715

**Authors:** G Kaur, T Lamb, A Tjitropranoto, Irfan Rahman

## Abstract

E-cigarettes (e-cigs) are a public health concern for young adults due to their rising popularity despite evidence of harmful effects. Yet an extensive study defining the cell-specific immune changes upon exposure to flavored e-cigs remains elusive. To determine the immunological lung landscape upon acute nose-only exposure of C57BL/6J to flavored e-cig aerosols, we performed single-cell RNA sequencing (scRNA seq). Analyses of the levels of metals in the e-cig aerosol generated daily during exposure revealed a flavor-dependent variation in the day-to-day leaching of the levels of metals like Ni, Cu, K and Zn, among others. scRNA profiles of 71,725 cells generated from control and treatment groups (n=2/sex/group) found maximum dysregulation of (a) myeloid cell function in menthol (324 differentially enriched genes (DEG)) and tobacco (553 DEGs) –flavor exposed and (b) lymphoid cell function in fruit (112 DEGs)-flavored e-cig aerosol exposed mouse lungs as compared to air. Flow cytometry analyses identified marked increase in the neutrophil percentage and a decrease in the eosinophil count in menthol and tobacco-flavored e-cig aerosol exposed mouse lungs which corroborated with our scRNA seq data. We further found an: (a) increase in CD8+ T cell percentages, (b) upregulation of inflammatory genes like *Stat4*, *Il1bos*, *Il1b*, *Il1ra* and *Cxcl3* and (c) enrichment of terms like ‘T-helper cell 1cytokine function’ and ‘NK cell degranulation’ in the lungs of e-cig aerosol exposed mouse when compared to control. Interestingly, the increase in the level of immature neutrophils characterized by Ly6G deficiency and reduction in the S100A8 (marker for neutrophil activation) using immunofluorescence in tobacco-flavored e-cig aerosol exposed mouse lung sections point towards a possible shift in the neutrophil dynamics upon exposure to e-cigs. Overall, this study identifies flavor dependent changes in myeloid and lymphoid-cell mediated responses in the mouse lungs exposed to acute nose-only e-cig aerosols and provides a resource to influence future research in select /specific cell types to understand the immunological implications of long-term use of e-cigs.

## Introduction

Electronic cigarettes (e-cigs) or electronic nicotine delivery systems are a relatively novel set of tobacco/nicotine and flavored products that have gained immense popularity amongst adolescents and young adults in many countries including the United States (US), the United Kingdom (UK) and China. Flavors are one of the key features that make these products alluring to the younger diaspora (1). Reports indicate that in 2020 about 22.5% of high school students and 9.4% of middle school students in the US were daily vapers or e-cig users with fruit (66%), mint (57.5%), and menthol (44.5%) being the most commonly used flavors (2). However, not much is known about the flavor-specific effects of e-cig vaping on the health and immunity of an individual specially focusing on cell-types and gene transcripts.

E-cig products and aerosols are known to contain harmful constituents including formaldehyde, benzaldehyde, acrolein, n-nitrosamines, volatile organic compounds (VOCs), ketenes, and metal ions (3–5). Studies have indicated that exposure to e-cigs may enhance inflammatory responses, oxidative stress, and genomic instability in exposed cells or animal systems (6–8). Risk assessment (systemic) of inhaled diacetyl, a potential component of e-liquids, has estimated the non-carcinogenic hazard quotient to be greater than 1 amongst teens (9). Furthermore, clinical and *in vivo* studies have suggested that exposure to e-cig aerosols could impair innate immune responses in the host thus making them more susceptible to bacterial/viral infections. The bacterial clearance, mucous production, and phagocytic responses in these individuals are shown to be affected upon use of e-cigs (10–14).

However, cell-specific changes within the lung upon vaping is not fully understood, making it hard to determine the long-term health impacts of the use of these novel products. In this respect, single cell technology is a powerful tool to analyze gene expression changes within cell populations to study cellular heterogeneity and function (15–17). Such an investigation is important to deduce the health effects of acute and chronic use of e-cigs in young adults. **In this study**, we aim to determine the effects of e-cig exposure on mouse lungs at single cell level. To do so, we exposed C57BL/6J mice to 5-day nose-only exposure to air, propylene glycol: vegetable glycerin (PG:VG), fruit-, menthol– and tobacco-flavored e-cig aerosols. The nose-only exposure has more translational relevance over the whole-body exposure (18), owing to which, we chose nose-only exposure profile for this work. To limit the stress to the animals a 1-hour exposure was chosen per day. We performed single-cell RNA sequencing (scRNA seq) on the lung digests from exposed and control animals and identified neutrophils and T-cells, among others, as the major cell populations in the lung that were affected upon acute exposure. We were able to identify 29 gene targets that were commonly dysregulated amongst all our treatment groups upon aggregating results from the major lung cell types. These gene targets are the markers of early immune dysfunction upon e-cig aerosol exposure *in vivo* and could be studied in detail to understand temporal changes in their expression and function that may govern allergic responses and adverse pulmonary health outcomes upon acute and sub-acute exposures to e-cigarette aerosols.

## Results

### Exposure to e-cig aerosols results in flavor-dependent exposure to different metals and mild histological changes in vivo

This study was designed to characterize the effects of exposure to flavored e-cig aerosols at single-cell level to understand the immunological changes in the lung microenvironment. To do so, we generated a single-cell profile of e-cig aerosol exposed mouse lungs (n= 2/sex/group). The thus obtained results were then validated with the help of our validation cohort of n= 3/sex/group as shown in **Figure 1A**. Since all the commercially available e-liquids used in this study contained tobacco derived nicotine (TDN), we first determined the levels of serum cotinine (a metabolite of nicotine) to prove successful exposure of the mice in each treatment group. As expected, we did not see any traces of cotinine in the serum of air and PG:VG exposed mice. Significant levels of cotinine were detected in the serum of mice exposed to fruit-, menthol-, and tobacco-flavored e-cig aerosols **(Figure S1B**), validating successful exposure of our test animals to TDN containing flavored e-cigs.

**Figure 1:**
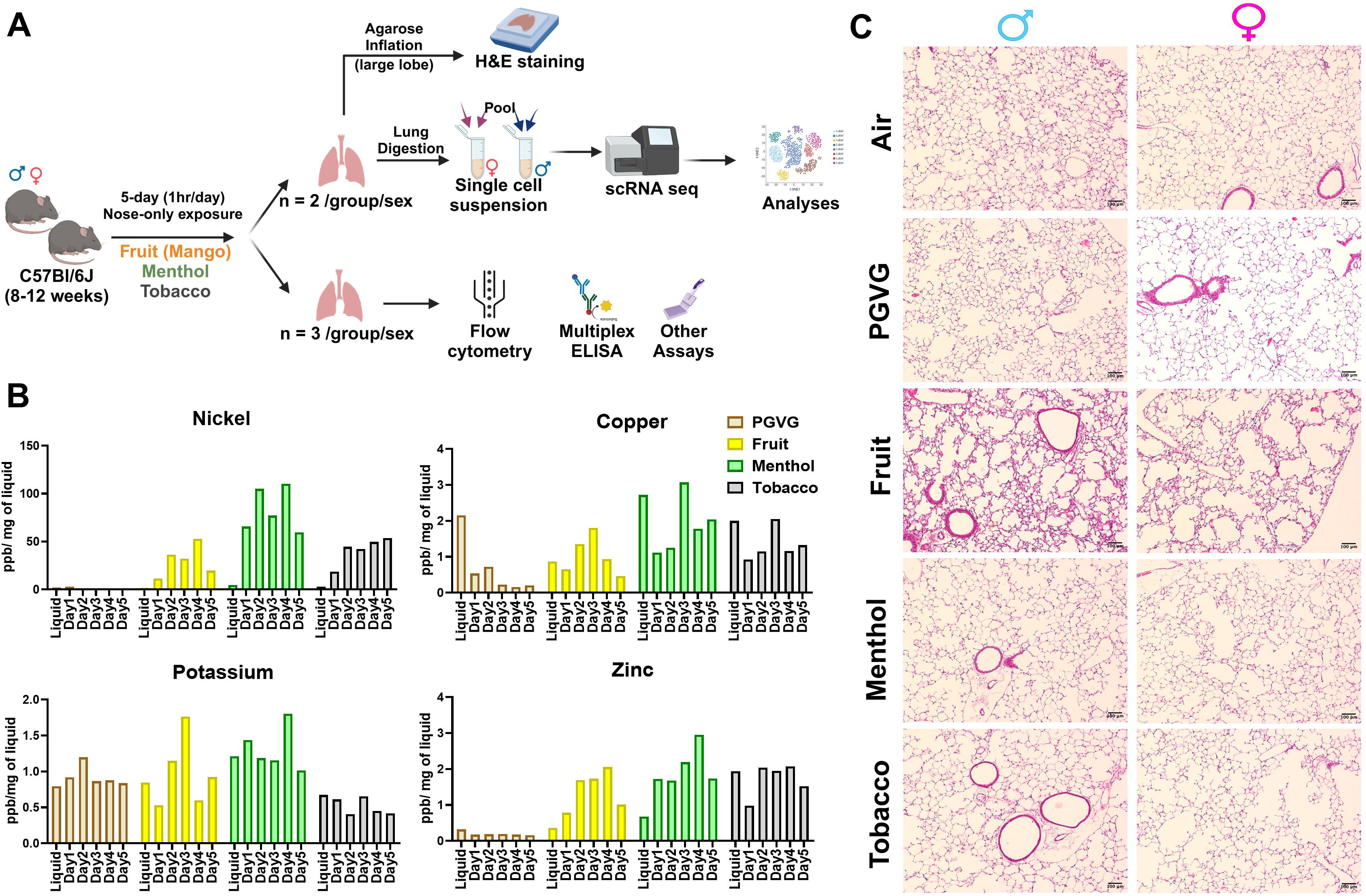
Flavor-dependent changes in the levels of quantified metals, but no major histological damage on acute exposure to flavored e-cig aerosol in C57BL/6J mice. Schematics showing the exposure profile and experimental design to understand the effects of exposure to differently flavored (fruit, menthol and tobacco) e-cig aerosols in the lungs of C57BL/6J mice using scRNA seq (A). Bar Graph showing the levels of metals as determined by ICP-MS in the aerosols captured daily during exposures using inExpose nose-only inhalation system from Scireq technologies (B). Lung morphometric changes observed using H&E staining of lung slices from air, PG:VG and differently flavored e-cig aerosol exposed mice lungs. Representative images of n = 2/sex/group at 10X magnification is provided (C).

Since metals released upon heating of the coils of e-cig devices are a source of toxicity upon vaping (19, 20), we further monitored the levels of metals in the e-cig aerosols generated during each day of mouse exposures. This acted as an indirect measure for characterizing the chemical properties of the aerosols used for exposure in this study. To monitor the release of metals into the mouse lungs, the aerosol condensate from each day of exposure was collected and the levels of select elements were detected using ICP-MS. A detailed account of the concentrations of identified elements/metals is provided in **Table 1**. Interestingly, we identified flavor-dependent changes in the levels of metals like Ni, Zn, Na, K and Cu on a day-to-day basis. Note, despite the use of same wattage and temperature (max of 230 C) for generation of e-cig aerosols, the leaching of each metal varied per day of exposure **(Figure 1B)**. This is a crucial result as it highlights the importance of studying the impact of atomizer, coil composition and design on the chemical composition of the generated aerosols. These variations might affect the risk and toxicity associated with each of these products, an area that has been recently explored by our group (21).

**Table 1:**
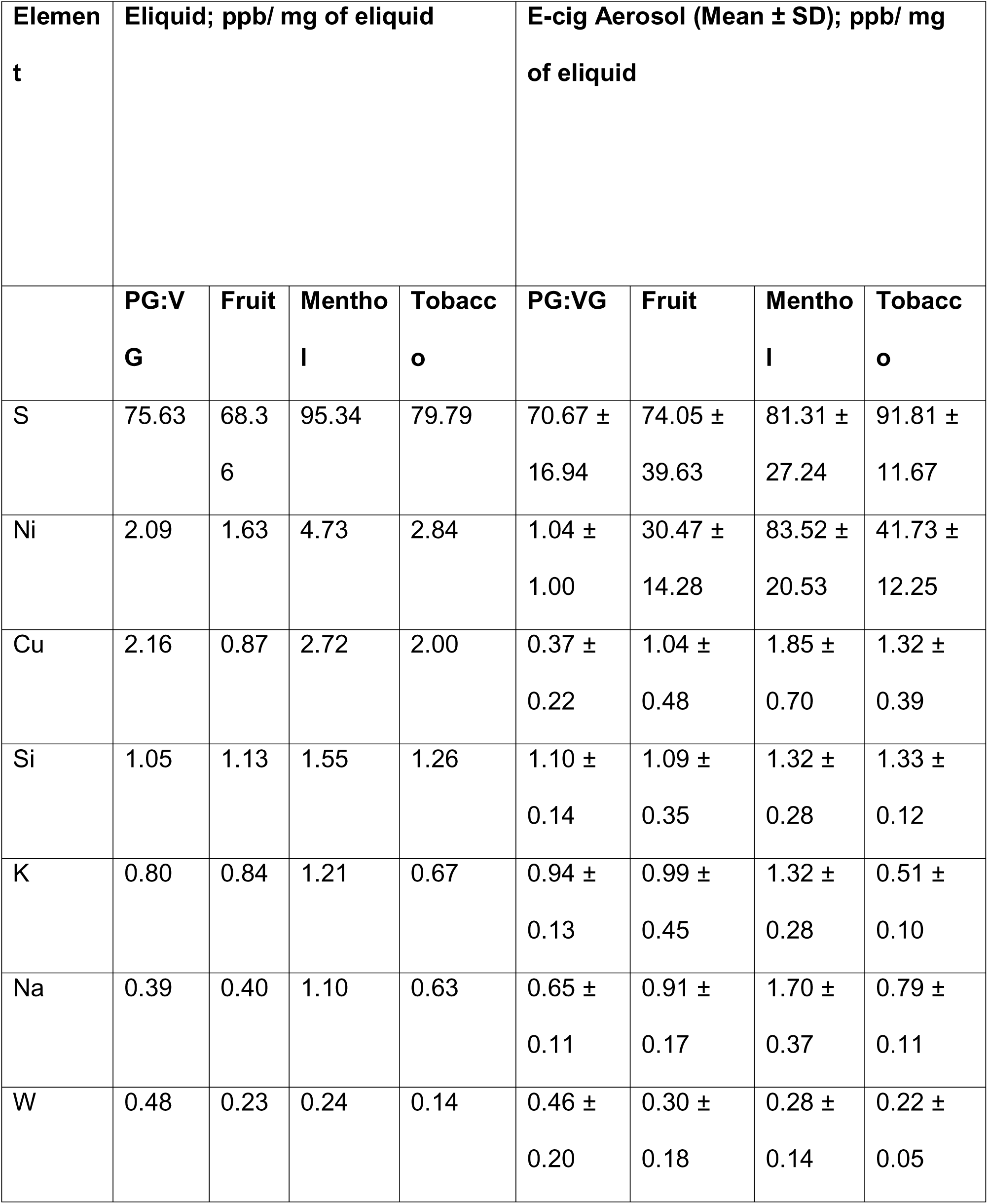
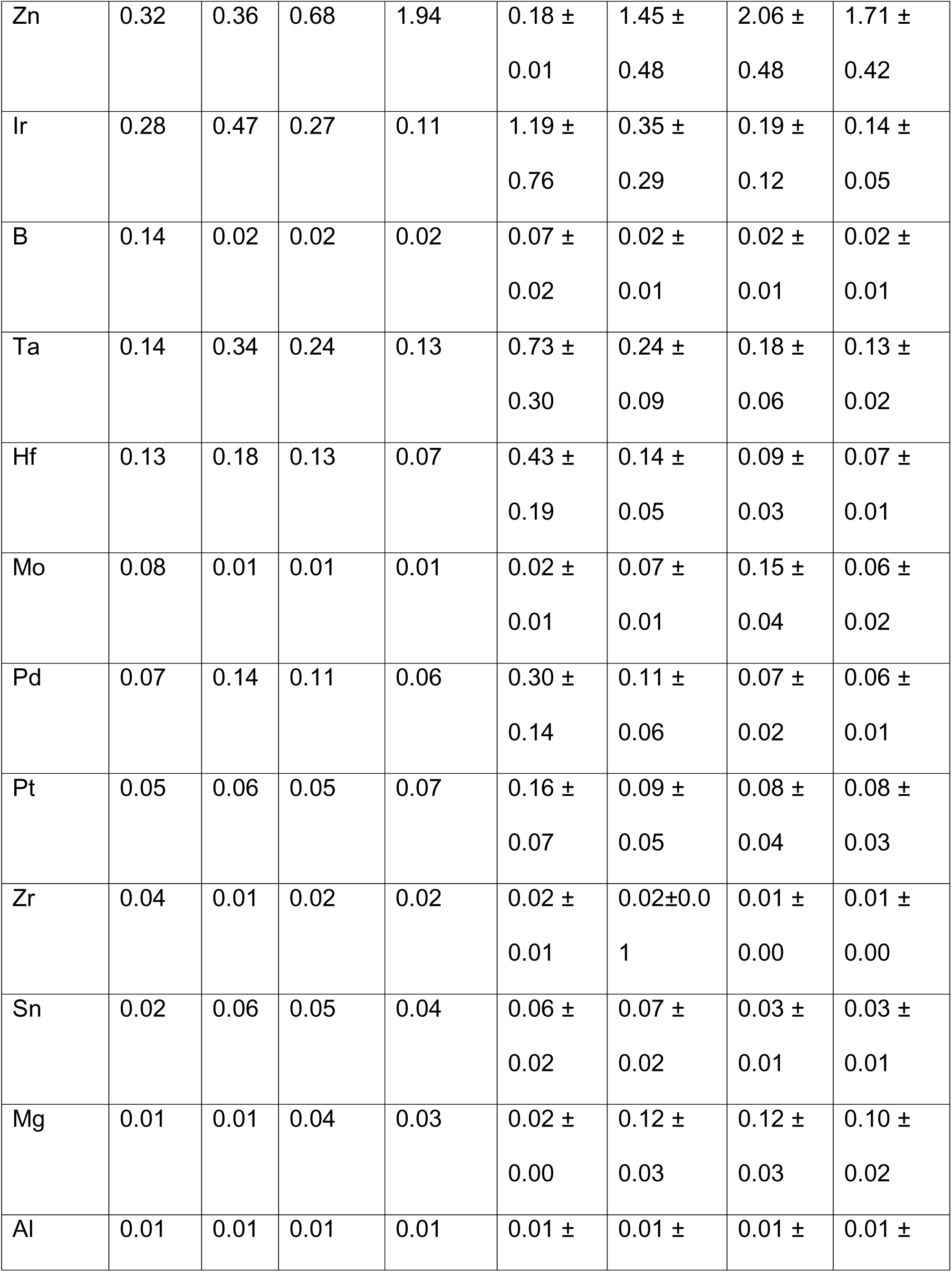
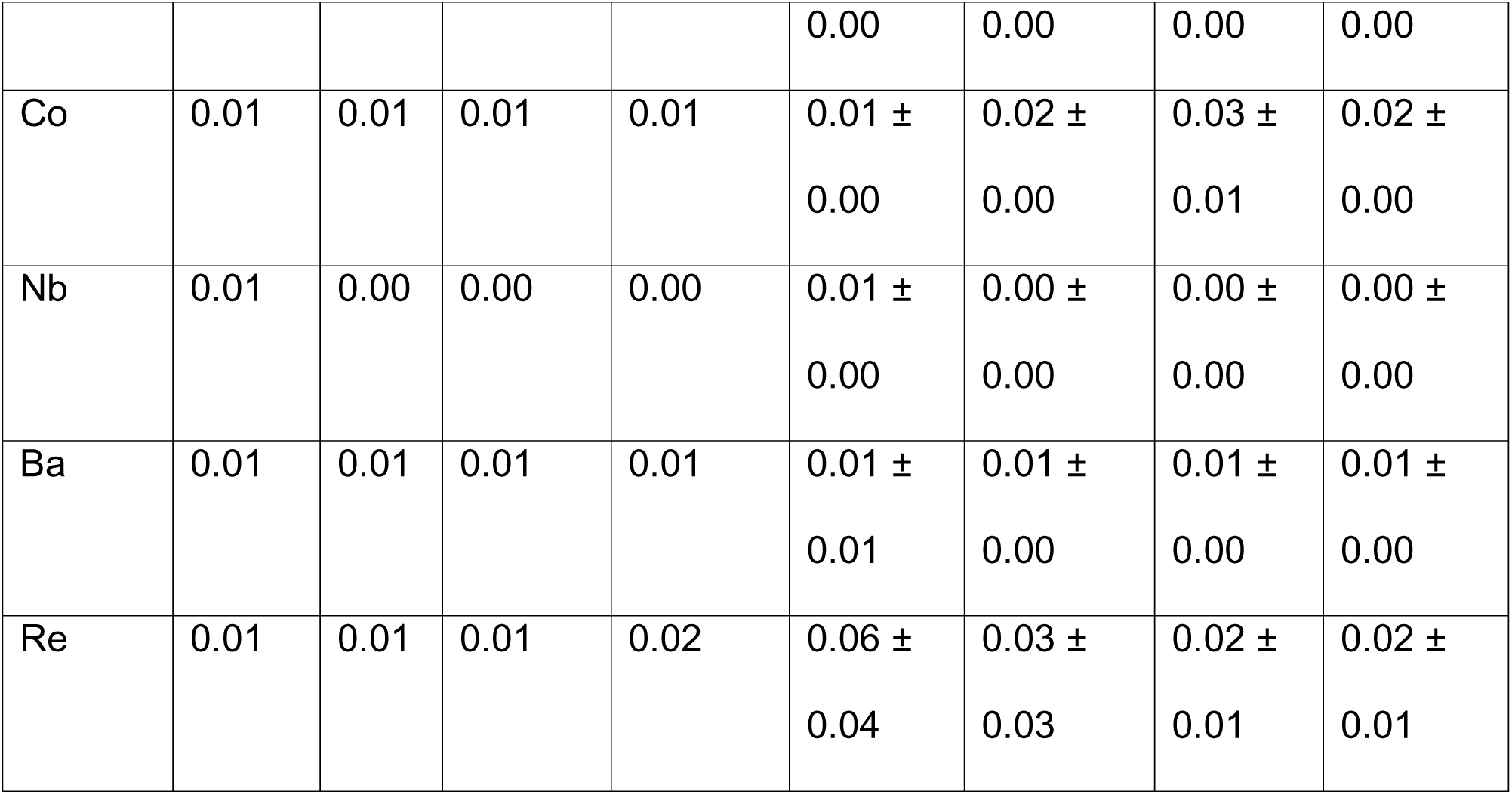
Table showing the levels of common elements found in the flavored e-liquids and e-cig aerosols as measured using ICP-MS.

Next, we performed H&E staining on the lung tissue sections to study the morphometric changes in the mouse lungs upon exposure to differently flavored e-cig aerosols. We did not find much evidence of tissue damage or airspace enlargement upon acute exposures in our model, as expected. However, we found evidence of increased alveolar septa thickening in the lungs of both male and female mice exposed to fruit-flavored e-cig aerosols **(Figure 1C)**. This could be a result of lower agarose inflation observed for the mouse lungs in this group. We had challenges/difficulties with inflating these mouse lungs. Owing to the lack of proper inflation, the lungs were in a collapsed state which could be a probable explanation for our histological observation. Since this was not the prime focus of our study, we did not conduct further experiments to confirm our speculations.

### Detailed map of cellular composition during acute exposure to e-cig aerosols reveals distinct changes in immune cell phenotypes

As the principle focus of this study was to identify the flavor-dependent and independent effects upon exposure to commercially available e-cig aerosols at the single-cell level, we performed the scRNA seq on the mouse lungs from exposed and control mice. After quality control filtering **(Figure S2),** normalization and scaling, we generated scRNA seq profiles of 71,725 cells in total. Except for the PG:VG group, all the rest of the treatments had approximately similar cell viabilities, cell capture, and other quality assessments. However, for normalization, equal features/ genes were used across all the groups for subsequent analyses. A detailed account of the cell number (single-cell capture) and gene features identified before and after filtering upon QC check of sequenced data is provided in **Supplementary File S1**.

Uniform Manifold Approximation and Projection (UMAP) was used for dimensionality reduction and visualization of cell clusters. Cell annotations were performed based on the established cell markers in Tabula Muris database and available published literature and we identified 24 distinct cell clusters as shown in **Figure 2A**. The general clustering of individual cell types based upon the commonly known cell markers was used to identify –**Endothelial** (identified by expression of *Cldn5*), **Epithelial** (identified by expression of Sftpa1), **Stromal** (identified by expression of *Col3a1*), and **Immune** (identified by expression of *Ptprc*) cell populations **(Figure 2B)**. The ‘FindVariableFeatures’ from Seurat was used to identify cell-to-cell variation between the identified clusters and the top 2000 variable genes identified in each cluster have been elaborated in **Supplementary File S2**. We observed minor variations in the cell frequencies as observed through scRNA seq analyses within cell types across different treatment groups **(Figure S3A)**. The largest proportion of cells was found in the endothelial cell cluster (43.87% of total per sample), followed by lymphoid (26.43% of total per sample) and myeloid (16.79% of total per sample) clusters. A detailed account of the two-way ANOVA statistics for the general clustering with cell types and treatment groups as independent variables is provided in **Supplementary File S3A**.

**Figure 2:**
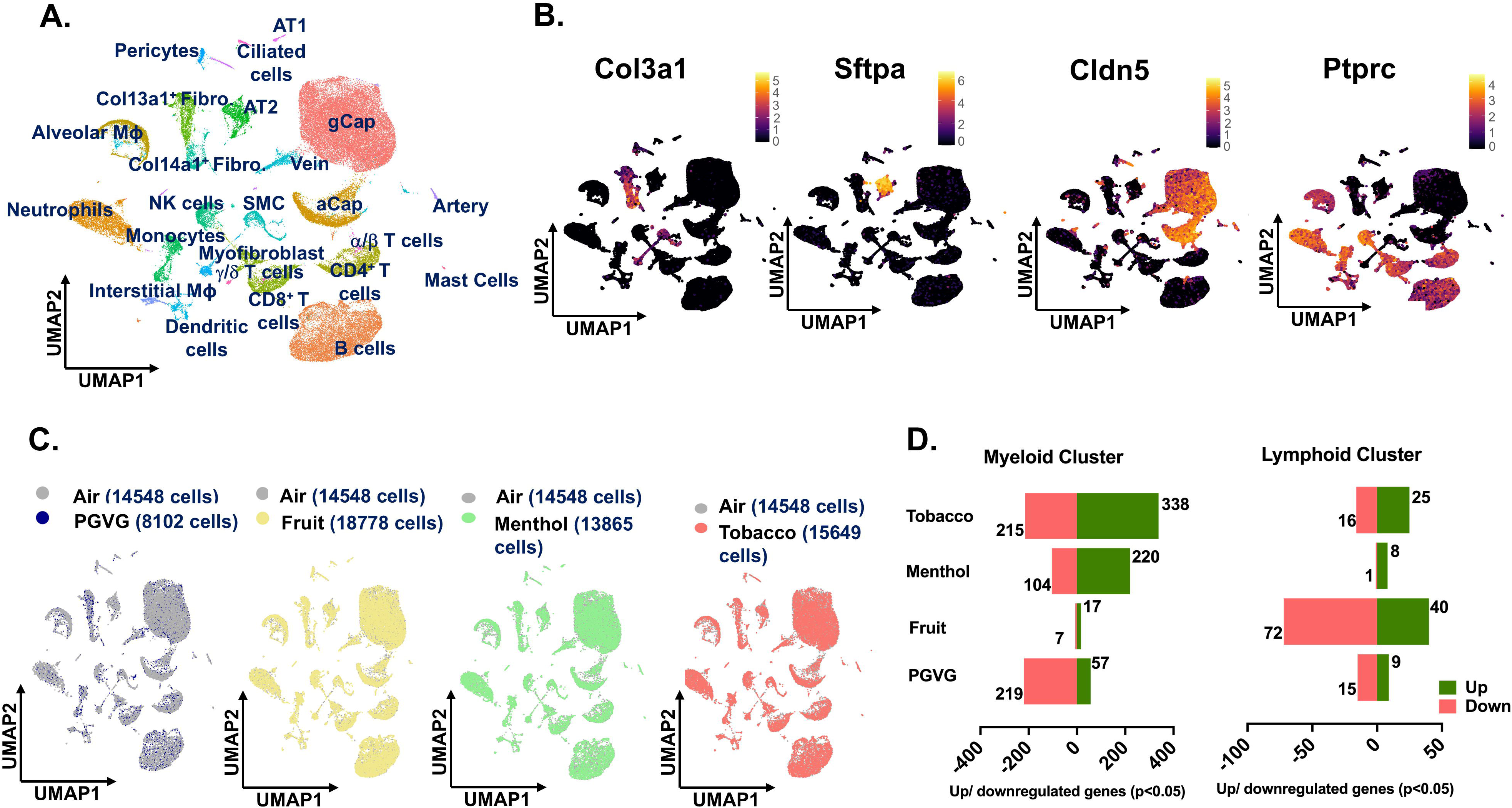
scRNA seq analyses reveal maximum changes in the transcriptional profile of immune cell population upon exposure to differently flavored e-cig aerosols. Male and female C57BL/6J mice (n = 2/sex/group) were exposed to 5-day nose-only exposure to differently flavored e-cig aerosols. The mice were sacrificed after the final exposure and lungs from air (control) and differently flavored e-cig aerosol (fruit, menthol, and tobacco)-exposed mice were used to perform scRNA seq. UMAP plot of 71,725 cells captured during scRNA seq showing the 24 major cell clusters identified from control and experimental mouse lungs **(A)** and the expression of canonical markers used for identifying stromal (*Col3a1*), epithelial (*S*ftpa1), endothelial (*Cldn5*) and immune (*Ptprc*) cell populations. The intensity of expression is indicated by the black-yellow coloring **(B)**. Group-wise comparison of the UMAPs upon comparing PG:VG (blue), fruit (yellow), menthol (green) and tobacco (red) versus air (grey) groups following dimensionality reduction and clustering of scRNA seq data **(C)**. Bar plot showing the number of significant (p < 0.05) differentially up-(green) and downregulated (red) genes in myeloid and lymphoid clusters in PGVG, fruit-, menthol-and tobacco-flavored e-cig aerosol exposed mouse lungs as compared to air controls. Here, AT1: alveolar type I, AT2: alveolar type II, Fibro: Fibroblast, MΦ: macrophage, SMC: smooth muscle cell, gCap: general capillary, aCap: alveolar capillary, and NK: natural killer.

Groupwise comparisons of each treatment group versus air control, did not show many changes in the cell clusters thus showcasing little to no effect on the overall lung cellular compositions in treated and control groups. However, the number of cells for PG:VG group was very low (8102 cells) as compared to other treatments (∼15,710 cells on average). We would like to report that this is an outcome of the low viability of this sample prior to scRNA seq, and not necessarily the effect of the treatment **(Figure 2C)**. Differential gene expression analyses showed dysregulation of genes in all cell populations, but maximum effect was observed on the cells from immune cell clusters. This is not surprising, as the immune system, especially the myeloid cells form the frontline of the host’s defenses against external stressors (22, 23). Compared to air, we observed a dysregulation of 553 genes (338 upregulated; 215 downregulated) in the myeloid cell cluster of mouse lungs exposed to tobacco-flavored e-cig aerosol. We identified 324 and 24 DEGs in the myeloid lung cluster from mouse lungs exposed to menthol– and fruit-flavored e-cig aerosols respectively as compared to air control **(Figure 2D; Supplementary Files S5-S10)**. For the lymphoid cluster, we observed maximum dysregulation in the lungs exposed to fruit-flavored e-cig aerosols with a total of 112 DEGs. In contrast, 41 and 9 significant DEGs were identified for lymphoid cluster from lungs exposed to tobacco and menthol-flavored aerosols respectively **(Figure 2D; Supplementary Files S12-S17**).

It is important to mention here that most of the commercially available e-liquids /e-cig products use PG:VG as the base liquid to generate the aerosol and act as a carrier for flavoring chemicals. Thus, we compared the effect of PG:VG alone in our study to make further comparisons between individual flavoring products with that of PG:VG only. DESeq2 analyses showed dysregulation of 276 genes in mouse lungs exposed to PG:VG alone in the myeloid cell cluster as compared to air controls **(Figure 2D; Supplementary Files S4)**. Contrary to this, exposure to PG:VG aerosols affected 24 genes in the lymphoid cluster as compared to air control **(Figure 2D; Supplementary Files S11)**. Furthermore, when compared to PG:VG, exposure to fruit, menthol and tobacco-flavored e-cig aerosol dysregulated 262, 873 and 960 genes in the myeloid cluster and 37, 64 and 112 genes in the lymphoid cluster respectively. A detailed account of the DEGs identified upon comparing fruit, menthol and tobacco-flavored e-cig aerosol exposed mouse lungs to PG:VG in myeloid and lymphoid clusters have been provided in **Supplementary Files S19-S24.**

### Exposure to e-cig aerosols results in flavor-dependent changes in neutrophilic and eosinophilic response in vivo

To study the specific changes in the innate and adaptive immunity upon acute exposure to differently flavored e-cig aerosols, we first compared the changes in the overall cell population of individual cell types using scRNA seq. The scRNA seq data was validated with the help of flow cytometry using a larger cohort of animals. Since we observed sex-dependent variations in oxidative stress responses upon exposure to e-cig aerosols in previous studies (24–26), the flow cytometry data was analyzed in a sex-dependent fashion.

We did not observe any changes in the cell frequencies of alveolar macrophages across treatments through scRNA seq analyses. In general, there was a moderate decrease in the cell frequencies of alveolar macrophages in exposed mice as compared to air control independent of the flavor profile of the e-liquid employed **(Figure 3A)**. Flow cytometric analyses confirmed little to no change in the alveolar macrophage (CD45^+^ CD11b^−^ SiglecF^+^) percentages within the lung of exposed versus control mice **(Figure 3C-D; Supplementary File S3B)**. Contrarily, we observed a flavor-dependent moderate increase in the cell frequencies of neutrophil cluster in menthol (0.1 ± 0.02) %– and tobacco (0.1 ± 0.06) %-flavored aerosol exposed mouse lung as compared to controls (0.06 ± 0.02) % **(Figure 3B)**. We used flow cytometry to validate the scRNA seq changes. Flow cytometry analyses showed an increase in the neutrophil (CD45^+^ CD11b^+^ Ly6G^+^) percentages of menthol-flavored e-cig aerosol exposed mouse lung corroborating with the scRNA seq results. Furthermore, our results showed this increase to be more pronounced in male mice (p = 0.0880) as compared to their female (p = 0.9662) counterparts **(Figure 3C-D, Supplementary File S3B).**

**Figure 3:**
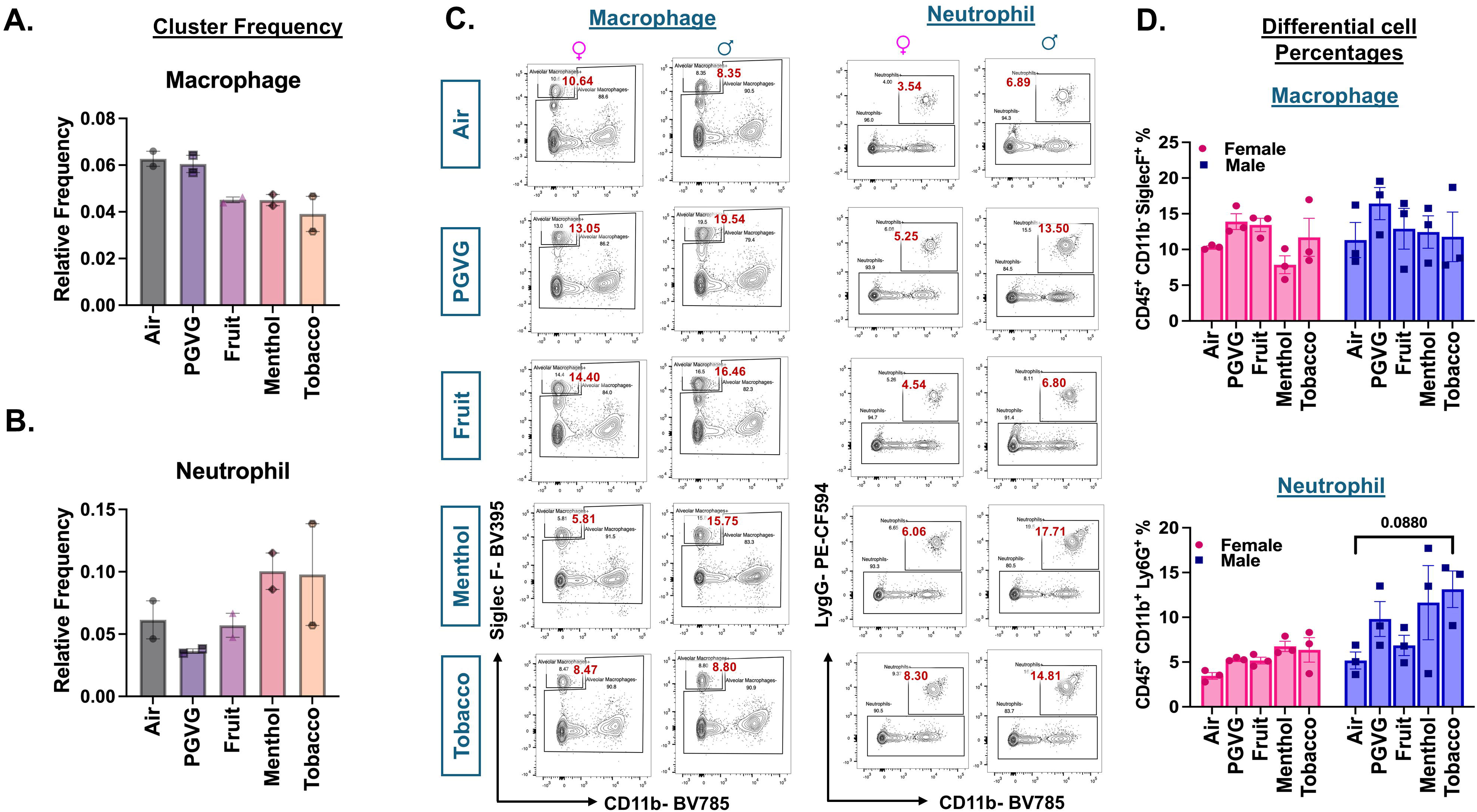
Cellular composition of myeloid cells in air and e-cig aerosol exposed mouse lungs reveal increase in neutrophil count through scRNA seq and flow cytometry. Relative cell frequencies of alveolar macrophages **(A)** and neutrophils **(B**) across controls and flavored e-cig aerosol exposed mouse lungs as determined using scRNA seq. Representative flow plots **(C)** and bar graphs **(D)** showing sex-dependent changes in the percentages of neutrophils (CD45+ CD11b+ Ly6G+) and alveolar macrophage (CD45+ CD11b-SiglecF+) populations in lung digests from mice exposed to differently flavored e-cig aerosols. Values plotted and written in red on the flow plots are representative of the percentage of each cell population in the total CD45+ cells present in the lung homogenates from treatment and control groups. Data are shown as mean ± SEM (n = 3/sex/group). SE determined using two-way ANOVA with a Tukey post-hoc test for all cell means, to analyze the main effects of sex and treatment and their interaction. The two-way ANOVA results are shown in **Supplementary File S3B.**

It is important to mention that due to the presence of nicotine in all the e-liquids used as treatments for this study, we added an extra control group-PG:VG+Nic-for select experiments. Flow cytometric analyses revealed slight variations in the lung neutrophils and macrophage percentages observed in PG:VG+Nic group as compared to PG:VG only. However, none of these changes were significant. Furthermore, the patterns of change observed for the lungs exposed to aerosols from flavored e-liquids were quite distinct from those observed for PG:VG+Nic thus proving that the observed changes are not solely due to the presence of nicotine in these groups **(Figures S3B)**.

Though we did not find a distinct eosinophil cluster in our scRNA seq analyzed data, interesting sex– and flavor-specific changes were observed in the lung eosinophil (CD45^+^ CD11b^+/-^ CD11c^−^ Ly6G^−^ SiglecF^+^) population upon analyzing the flow cytometry results. A significant decline in the eosinophil percentage was observed in the lung homogenates from menthol (p =0.0113) and tobacco (p = 0.0312)-flavored e-cig aerosol exposed male C57BL/6J mice as compared to air control. Contrarily, the eosinophil levels in the lung of male mice exposed to fruit flavored e-cig aerosols did not show remarkable changes when compared to the levels found in control **(Figure 4 & S4A)**. These results further emphasize the need of studying the sex-specific changes in the immune responses upon exposure to e-cig aerosols *in vivo*.

**Figure 4:**
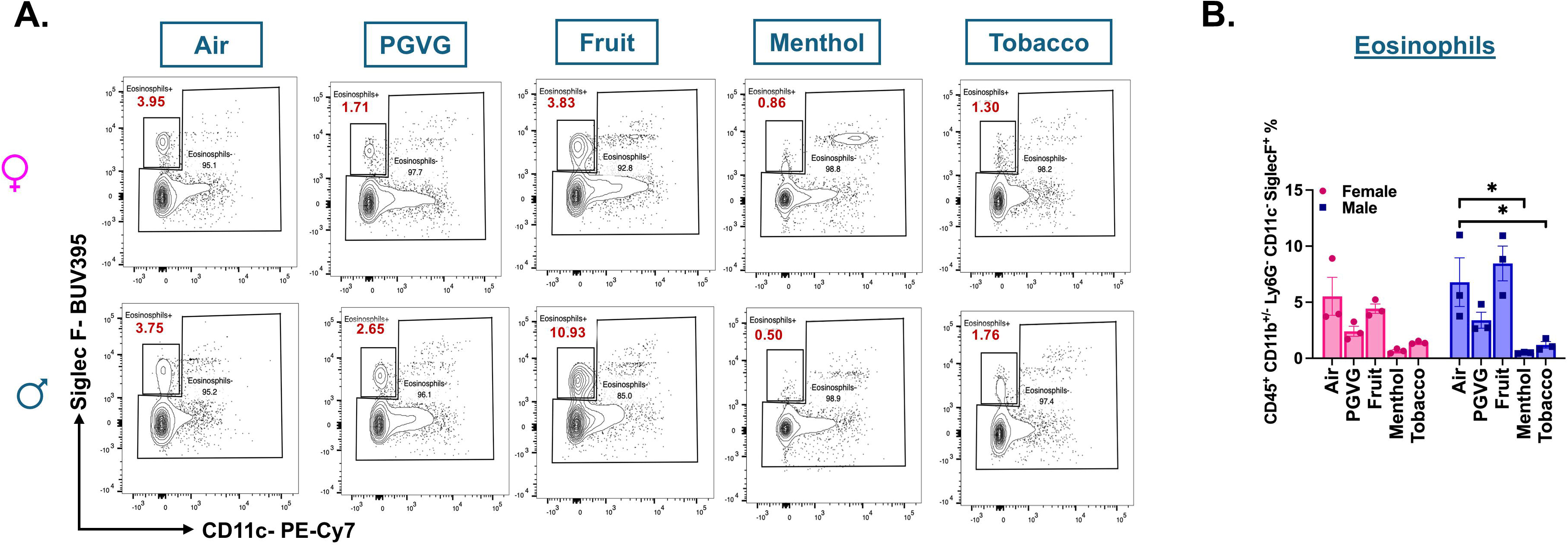
Flow cytometry analyses show significant decrease in the percentage of eosinophils in the lungs of menthol and tobacco-flavored e-cig aerosol exposed C57BL/6J mice. Representative flow plot **(A)** and bar graphs **(B)** showing the changes in the percentages of eosinophils (CD45^+^ CD11b^+/-^ CD11c^−^ Ly6G^−^ SiglecF^+^) found in the lungs of differently flavored e-cig aerosol exposed mouse lungs as compared to air controls. Data are shown as mean ± SEM (n = 3/sex/group). *p<0.05, per two-way ANOVA with a Tukey post-hoc test for all cell means, to analyze the main effects of sex and treatment and their interaction. The two-way ANOVA results are shown in **Supplementary File S3B**. Values plotted and written in red on the flow plots are representative of the percentage of each cell population in the total CD45+ cells present in the lung homogenates from treatment and control groups.

### Activation of T-cell cytotoxic responses in lymphoid cells upon exposure to tobacco-flavored e-cig aerosols

We next studied the changes in the lymphoid clusters of treated and control samples. While we did not notice any change in the CD4+ T cell frequencies in treatment and control groups using scRNA seq, our flow cytometry data showed significant increase in the CD4+ T-cell (CD45^+^ CD11c^−^ Ly6G^−^ CD11b^−^ MHCII^−^ CD4^+^) frequencies in the lungs of tobacco-flavored e-cig aerosol exposed female (p= 0.0492) C57BL/6J mice **(Figures 5A, 5C-D).** scRNA data did not show any change in the cell frequencies of CD8+ T cells in control and exposed mouse lungs either **(Figures 5B)**. However, flow cytometric analyses presented a different picture. We found a significant sex-independent increase in the CD8+ T (CD45^+^ CD11c^−^ Ly6G^−^ CD11b^−^MHCII^−^CD8^+^) cell percentages in the lungs of menthol– and tobacco-flavored e-cig aerosol exposed mice as compared to air control. Contrarily, the CD8+ T cell percentages increased in the fruit-flavored e-cig aerosol exposed male (p= 0.0163) mouse lungs and not in their female counterparts **(Figure 5C-D, Supplementary File S3B)**. Of note, we did not observe any changes in the CD4+ and CD8+ T cell percentages in PGVG+Nic group as compared to control, reiterating that the e-cig flavors are responsible for altered immune responses upon acute exposure in mice **(Figure S4B-C)**.

**Figure 5:**
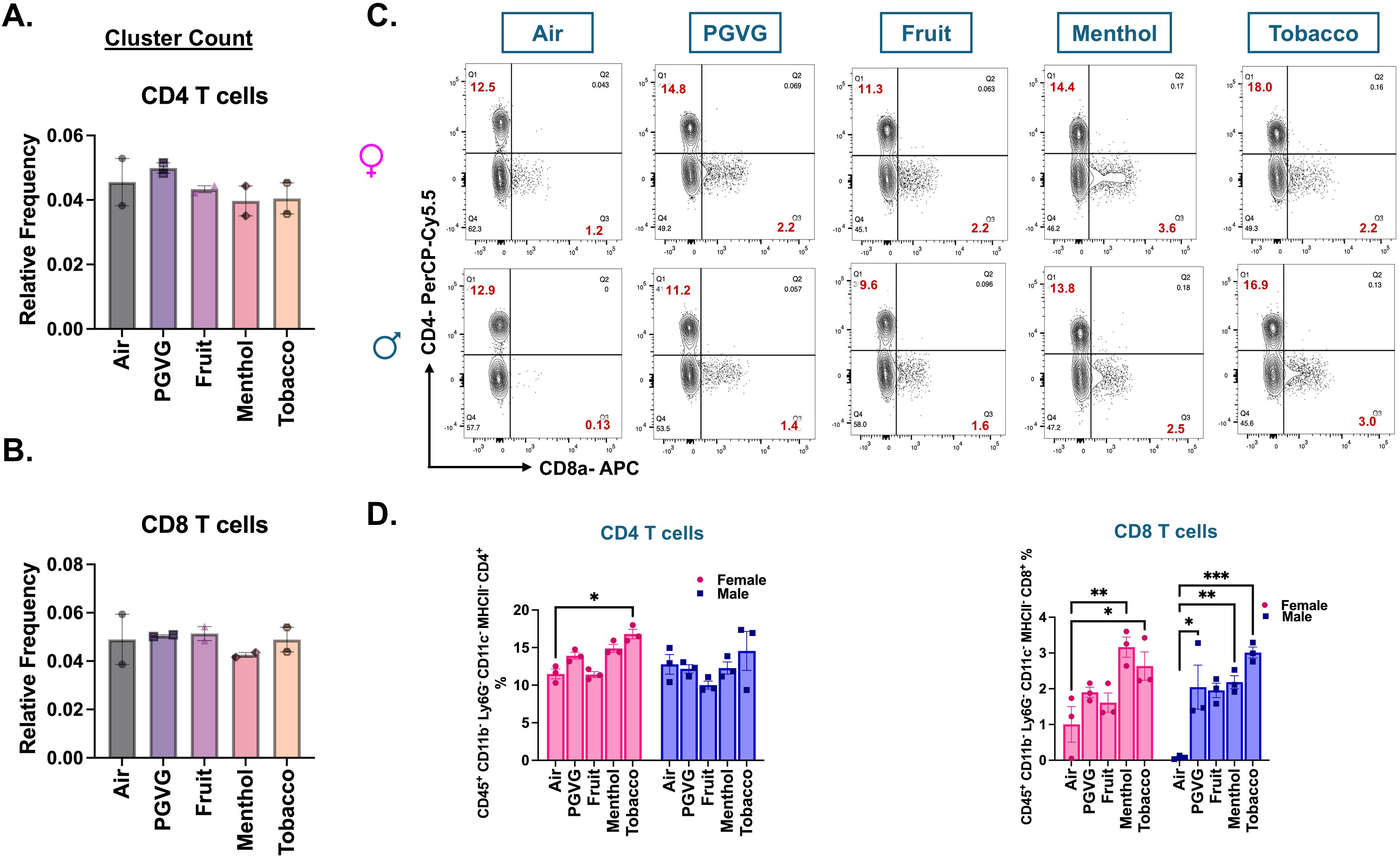
Flow cytometry results show sex-specific flavor-dependent increase in CD8^+^ T cells in lungs of differently flavored e-cig aerosol exposed C57BL/6J mouse. Relative cell frequencies of CD4^+^ **(A)** and CD8^+^ **(B)** T-cells across controls and flavored e-cig aerosol exposed mouse lungs as determined using scRNA seq. Representative flow plots **(C)** and bar graph **(D)** showing changes in the mean cell percentages of CD4^+^ (CD45^+^ CD11c^−^ Ly6G^−^ CD11b^−^ MHCII^−^ CD4^+^) and CD8^+^ (CD45^+^ CD11c^−^ Ly6G^−^ CD11b^−^ MHCII^−^ CD8^+^) T-cells in the lung tissue digests from male and female mice exposed to differently flavored e-cig aerosols as determined using flow cytometry. Values plotted and written in red on the flow plots are representative of the percentage of each cell population in the total CD45+ cells present in the lung homogenates from treatment and control groups. Data are shown as mean ± SEM (n = 3/sex/group). *p<0.05, **p<0.01 and ***p<0.001; per two-way ANOVA with a Tukey post-hoc test for all cell means, to analyze the main effects of sex and treatment and their interaction. The two-way ANOVA results are shown in **Supplementary File S3B.**

### Sub-clustering of myeloid cluster identifies a population of neutrophils devoid of Ly6G

To probe further, we subclustered the myeloid cell populations. Upon sub clustering, we identified 14 unique clusters comprising all the major cell phenotypes including neutrophils, alveolar macrophages (AM), interstitial macrophages (IM), monocytes, dendritic cells (DC), and mast cells (**Figure 6A)**. A detailed account of all the cell types identified with their respective marker genes in myeloid subcluster is provided in **Supplementary File S18**. On deeper evaluation, we identified two unique phenotypes of neutrophils (identified by *S100a8*, *S100a9*, Il1b, *Retnlg, Mmp9* and *Lcn2)* in the mouse lungs. These clusters were named as ‘Ly6G^+^ Neutrophils’ and ‘Ly6G^−^ Neutrophils’ based on the presence or absence of Ly6G marker, respectively **(Figure 6B)**. Ly6G is important for neutrophil migration, maturation and function within the lung (27). Since we did not anticipate identifying such a population of neutrophils following acute e-cig exposure, we did not gather flow cytometry evidence to validate this finding. We speculated that the Ly6G-population is a population of immature neutrophils. To confirm this possibility, we performed co-immunofluorescence using S100A8 (red; marker for neutrophil activation) and Ly6G (green) specific antibodies for control (air) and tobacco-flavored e-cig aerosol exposed mouse lungs. Co-immunofluorescence results showed a moderate increase (p =0.4429) in the level of Ly6G+ cells in the lungs of tobacco-flavored aerosol exposed mice as compared to air. However, the level of S100A8+ cells decreased markedly (p = 0.0571) in e-cig exposed group, thus showing that activation of neutrophils could be affected upon acute exposure to e-cigs **(Figures 6C-D; Supplementary Figure S4D)**.

**Figure 6:**
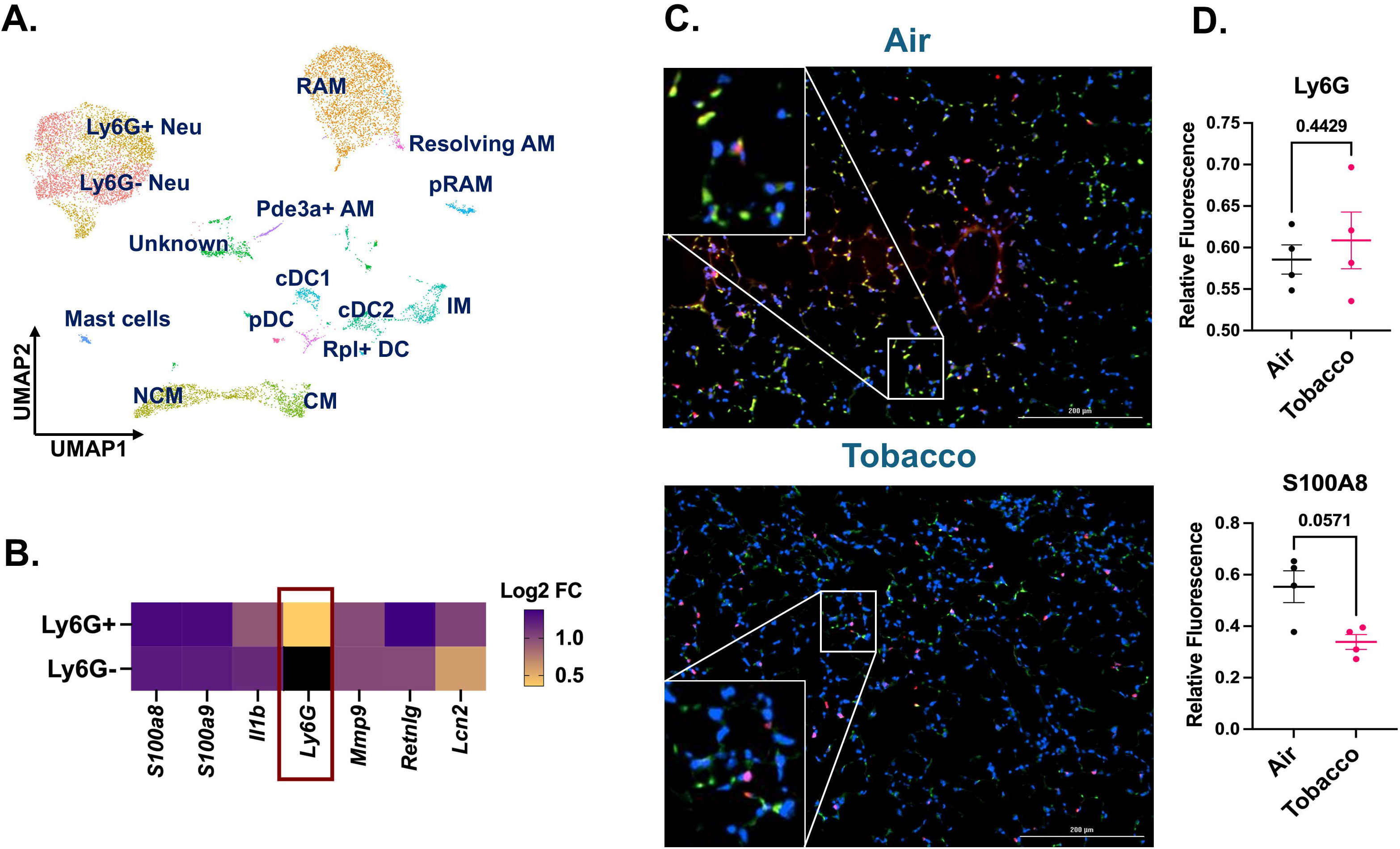
Co-immunofluorescence validates the increase of Ly6G– and Ly6G+ neutrophil population in the tobacco flavored e-cig exposed female C57BL/6J mice. The myeloid cell clusters were subsetted to identify two populations of neutrophils with and without the presence of Ly6G cell marker representing mature and immature neutrophils respectively. UMAP showing the 14 distinct cell populations identified upon subsetting and re-clustering the myeloid clusters from scRNA seq dataset from control and e-cig exposed mouse lungs **(A)**. Marker plot showing the differential expression of highly expressed genes in the Ly6G+ and Ly6G-neutrophil cluster. The intensity of expression is indicated by the yellow-blue coloring; black represents nil value for expression for that gene **(B)**. scRNA seq findings for presence of mature (Ly6G+) and immature (Ly6G-) neutrophils were validated by staining the tissue sections from tobacco-flavored e-cig aerosols and control (air) with Ly6G (green) and S100A8 (red, neutrophil activation marker). Representative images showing the co-immunostaining of Ly6G and S100A8 (shown as yellow puncta) **(C)** with respective quantification of relative fluorescence for Ly6G and S100A8 **(D)** in control and tobacco-flavored e-cig aerosol exposed mice. Data are shown as mean ± SEM (n = 4/group). SE calculated per Mann-Whitney U test for pairwise comparisons. Here, Neu: neutrophil, AM: alveolar macrophage, RAM: resident AM, pRAM: Proliferating RAM, IM: interstitial macrophage, CM: classical monocyte, NCM: non-classical monocyte, and DC: dendritic cell.

### Exposure to fruit-flavored e-cig aerosols affect the mitotic pathway genes in the lymphoid cluster

To assess the effects of acute exposures to differently flavored e-cig aerosols on the cell composition in the mouse lungs, we performed differential expression analyses for each flavor in comparison to air and PG:VG controls. The fruit-flavored e-cig aerosol exposure had the mildest effect on the cell compositions and gene expression as compared to the controls in our study. We did not observe major dysregulation in the gene expression for myeloid cell cluster in fruit-flavored e-cig exposed mouse lungs as compared to air controls. A total of 24 genes (17 upregulated; 7 downregulated) were significantly (p < 0.05) differentially expressed in the myeloid clusters in the lungs of animals exposed to fruit-flavored e-cig aerosols **(Figure 7Ai, Supplementary File S5)**. GO analyses of differentially expressed (DE) up-regulated genes showed enrichment of terms like ‘NK cell mediated immune response to tumor cells’ (fold enrichment= 9.75; p_adj_ = 0.0003), ‘haptoglobin binding’ (fold enrichment= 7.76; p_adj_ = 0.0096) and ‘transmembrane-ephrin receptor activity’ (fold enrichment= 7.05; p_adj_ = 0.014) in **(Figures 7Aii, Supplementary File S6A-B**) fruit-flavored aerosol exposed mice lung as compared to control. Importantly, few gene clusters (*Hbb-bs*, *Hbb-bt*, *Hba-a2* and *Hbb-a1*) showed sex-specific changes in the gene expressions in exposed lungs as compared to control. These gene clusters enriched for ‘erythrocyte development’ and warrant further study.

**Figure 7:**
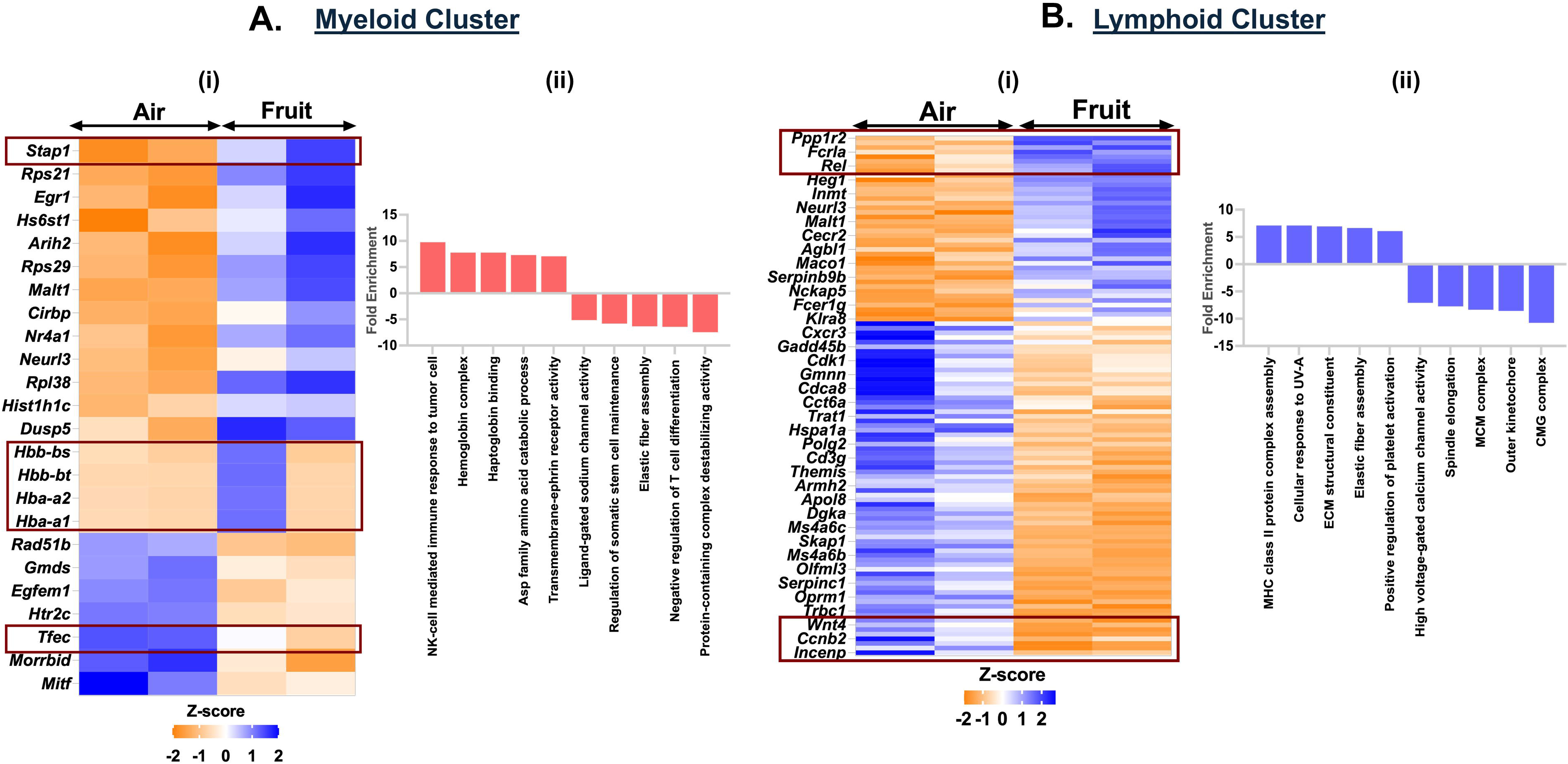
Exposure to fruit flavored e-cig aerosols result in activation of oxidative stress-mediated innate immunity in C57BL/6J mouse lungs. Male and female C57BL/6J mice were exposed to 5-day nose-only exposure to fruit-flavored e-cig aerosols. The mice were sacrificed after the final exposure and mouse lungs from air (control) and aerosol (fruit-flavored) exposed groups was used to perform scRNA seq. Heatmap and bar plot showing the DESeq2 **(i)** and GO analyses **(ii)** results from the significant (p<0.05) up/downregulated DEGs in the myeloid **(A)** and lymphoid **(B)** cell cluster from fruit-flavored e-cig aerosol exposed mouse lungs as compared to controls.

We further found dysregulation of 112 genes (40 up– and 72-down-regulated) in the lymphoid cluster of exposed mice. Upregulation of *H2-DMb2*, *H2-DMb1*, *Gp5*, *Pdpn* among others enriched for ‘MHC class II protein complex assembly’ (fold enrichment= 7.10; p_adj_ = 0.0009) and ‘positive regulation of platelet activation’ (fold enrichment= 6.08; p_adj_ = 0.0023) in exposed group as compared to air control. We also observed downregulation of genes like *Mcm4*, *Mcm2*, *Kif4*, *Cdca8*, *Cacna1f*, *Cacnb3*, in the lymphoid cluster of mice exposed to fruit-flavored e-cig aerosol. GO analyses of the downregulated genes enriched for terms like ‘CMG complex’ (fold enrichment= 10.75; p_adj_ = 3.39E-07), ‘spindle elongation’ (fold enrichment= 7.72; p_adj_ = 0.0005) and ‘high voltage-gated calcium channel activity’ (fold enrichment= 7.09; p_adj_ = 0.0059) in fruit-flavored aerosol exposed mouse lungs as compared to air controls**(Figure 7Bi-ii, Supplementary File S12-S13**).

### Exposure to Menthol-flavored e-cig aerosols affects the immune cell function

We demonstrated an upregulation of 220 genes and a downregulation of 104 genes in the myeloid cluster of mouse lungs exposed to menthol-flavored e-cig aerosol as compared to ambient air. We observed increased expression of inflammatory genes including *Il12b, Arid5a, Il12a, Il1b, Cacna1d and Cacnb2* enriching for terms like ‘T-helper 1 cell cytokine production’ (fold enrichment= 6.21; p_adj_ = 0.0115) and ‘L-type voltage-gated calcium channel complex’ (fold enrichment= 5.58; p_adj_ = 0.0191) in the myeloid cluster of menthol-flavored e-cig aerosol exposed mouse lungs as compared to air. We also found a downregulation in the expression of *Ovol1*, *Mapk15*, *Erbb2*, *Nrg4*, *Katnal2*, *Hspa1b* and *Hspa1a* in the myeloid cells of flavored e-cig aerosols. GO analyses of the DEGs, thus, showed enrichment of terms like ‘regulation of meiotic cell cycle phase transition’ (fold enrichment= 6.48; p_adj_ = 0.0051), ‘ERBB4 signaling pathway’ (fold enrichment= 5.40; p_adj_ = 0.025), and ‘protein-containing complex destabilizing activity’ (fold enrichment= 6.36; p_adj_ = 0.0095) as the top hits as shown in **Figure 8Ai-ii (Supplementary File S7-S8)**.

**Figure 8:**
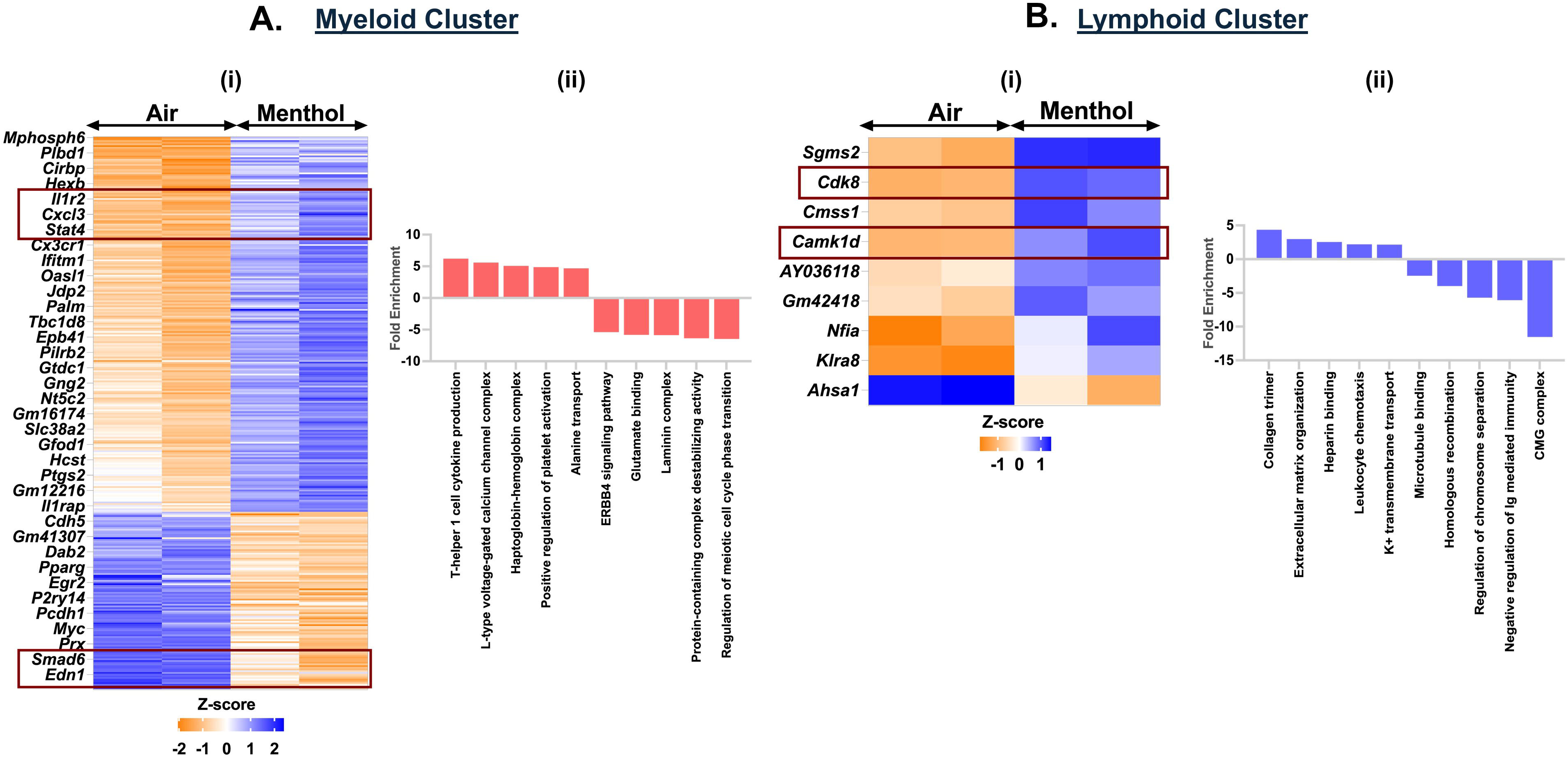
Exposure to Menthol flavored e-cig aerosols result in activation of innate immune responses C57BL/6J mouse lungs. Male and female C57BL/6J mice were exposed to 5-day nose-only exposure to menthol-flavored e-cig aerosols. The mice were sacrificed after the final exposure and mouse lungs from air (control) and aerosol (menthol-flavored) exposed groups was used to perform scRNA seq. Heatmap and bar plot showing the DESeq2 **(i)** and GO analyses **(ii)** results from the significant (p<0.05) up/downregulated DEGs in the myeloid **(A)** and lymphoid **(B)** cell cluster from menthol-flavored e-cig aerosol exposed mouse lungs as compared to controls.

Contrary to the responses observed for exposure to fruit-flavored e-cig aerosols, we found significant (p< 0.05) upregulation in the expression of *Cdk8* and *Camk1d* genes in the lymphoid cell population for menthol-flavored aerosol exposed mouse lungs **(Figure 8Bi, Supplementary File S14)**. *Cdk8* (cyclin-dependent kinase 8) is a transcriptional regulator that has a role in the cell cycle progression (28). Whereas *Camk1d* (calcium/calmodulin dependent protein kinase ID) functions to regulate calcium-mediated granulocyte function and respiratory burst within the cells(29). GO analyses of up and downregulated genes showed enrichment of terms including ‘regulation of toll-like receptor 9 signaling pathway’ (fold enrichment= 5.96; p_adj_ = 0.034), ‘transforming growth factor beta activation’ (fold enrichment= 5.42; p_adj_ = 0.044), ‘mitotic DNA replication’ (fold enrichment= –7.98.; p_adj_ = 2.51E-05) and ‘outer kinetochore’ (fold enrichment= –8.58.; p_adj_ = 1.13E-08) in the lymphoid cluster from menthol-flavored e-cig aerosol exposed mouse lungs **(Figure 8Bii, Supplementary File S15A-B)**.

### Exposure to Tobacco-flavored e-cig aerosol elicits immune response in myeloid cell and cell cycle arrest in lymphoid cell population

Like menthol-, tobacco-flavored e-cig aerosol also elicited a significant increase in the expression of 338 genes and a decrease in 215 genes as compared to air controls in the myeloid cell cluster. We observed an increase in the expression of chemokines like *Stat4*, *Il1b*, *Il1bos, Il18r1, Unc13d*, *Lgals9* and *Nkg7* in the myeloid cells resulting in enrichment of terms like ‘T-helper 1 cell cytokine production’ (fold enrichment= 6.66; p_adj_ = 0.0011) and ‘natural killer cell degranulation’ (fold enrichment= 6.28; p_adj_ = 0.0042) in tobacco-flavored e-cig aerosol exposed mouse lungs as compared to air controls.

We further found downregulation of genes including *Mcmdc2*, *Rad51*, *Spp1* and *Slc34a2* which enrich for terms like ‘double-strand break repair involved in meiotic recombination’ (fold enrichment= 6.44; p_adj_ = 0.0019) and ‘intracellular phosphate ion homeostasis’ (fold enrichment= 6.52; p_adj_ = 0.0052) in myeloid cluster from tobacco-exposed e-cig aerosols **(Figure 9Ai-ii, Supplementary File S9-S10)**.

**Figure 9:**
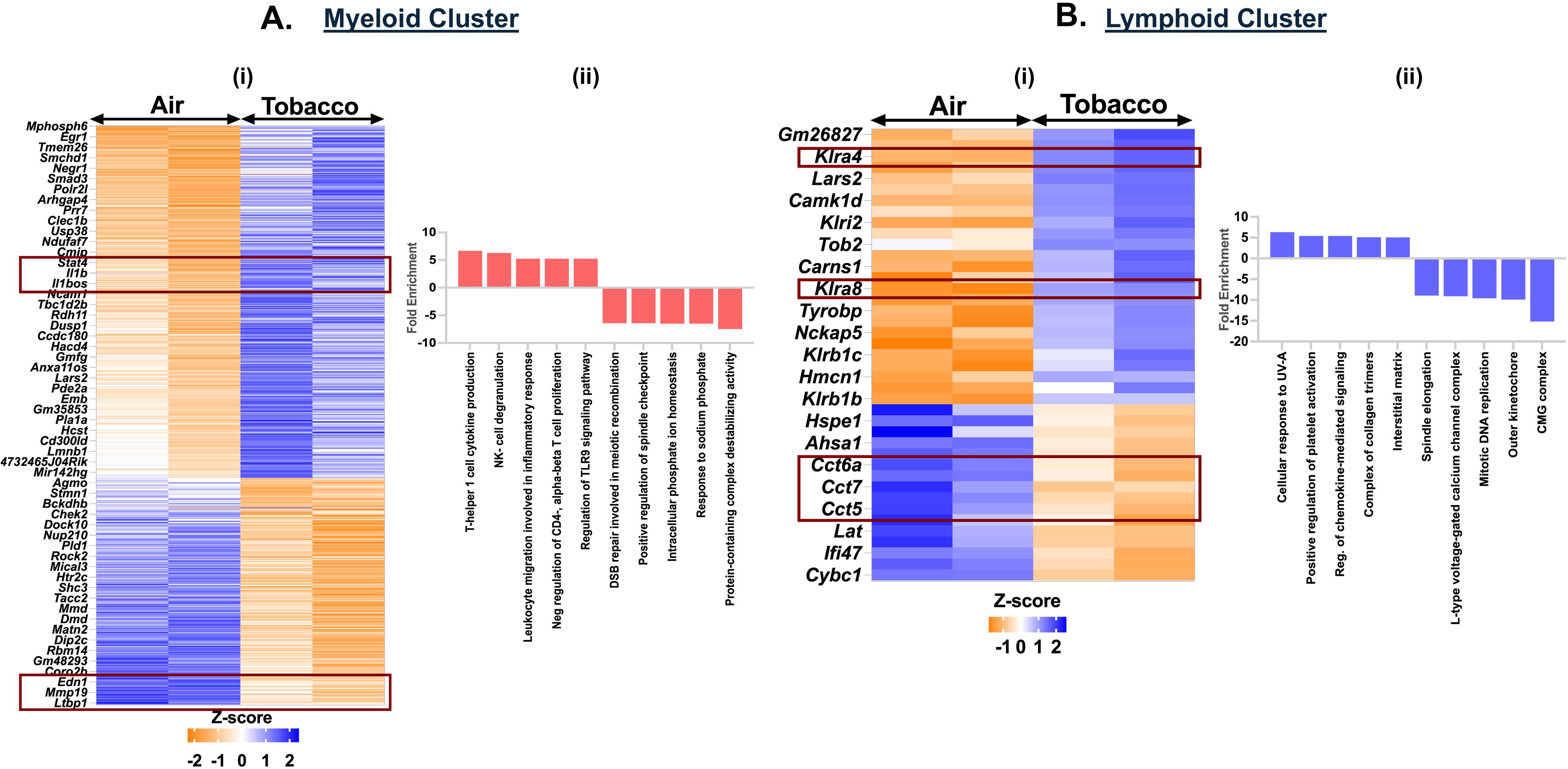
Exposure to Tobacco flavored e-cig aerosols result in activation of cytolysis and neutrophil chemotaxis in C57BL/6J mouse lungs. Male and female C57BL/6J mice were exposed to 5-day nose-only exposure to tobacco-flavored e-cig aerosols. The mice were sacrificed after the final exposure and mouse lungs from air (control) and aerosol (tobacco-flavored) exposed groups was used to perform scRNA seq. Heatmap and bar plot showing the DESeq2 **(i)** and GO analyses **(ii)** results from the significant (p<0.05) up/downregulated DEGs in the myeloid **(A)** and lymphoid **(B)** cell cluster from tobacco-flavored e-cig aerosol exposed mouse lungs as compared to controls.

We also observed a downregulation of genes responsible for chaperone-mediated protein folding (*Cct5*, *Cct7*, *Cct8*) in the lymphoid cluster from tobacco-flavored e-cig aerosol exposed mouse lungs. Downregulation of these genes could be indicative of the accumulation of misfolded proteins in these lungs which may lead to enhanced cell death (30, 31). DEG and GO analyses identified upregulation of *Robo1*, *Trem2*, *Padi2*, *Gp5*, *Gp9* and *Pla2g4a* and downregulation of *Mcm4*, *Mcm2*, *Rad51*, *Cacnb2*, and *Cacnb3* in the lymphoid cluster from tobacco-flavored e-cig aerosol exposed mouse lungs resulting in enrichment of terms including ‘regulation of chemokine-mediated signaling pathway’ (fold enrichment= 5.43; p_adj_ = 0.0304), ‘positive regulation of platelet activation’(fold enrichment= 5.43; p_adj_ = 0.0089), ‘mitotic DNA replication’ (fold enrichment= –9.6; p_adj_ = 6.63E-07), and ‘L-type voltage-gated calcium channel complex’ (fold enrichment= –9.12; p_adj_ = 0.0005) **(Figure 9Bi-ii, Supplementary File S16-S17)**.

### Dysregulation of chemokine signaling and T-cell activation on exposure to flavored e-cig aerosols

Since we showed increased production of cytokines/chemokines, driving the immune responses in mouse lungs exposed to flavored e-cigs, we performed multianalyte assay to determine the levels of these inflammatory cytokines in the lung digests from the exposed animals as shown in **Figure S4E**. Exposure to tobacco-flavored e-cig aerosol resulted in a marked increase in the levels of chemotactic chemokines including CXCL16, CXCL12, CXC3R and proinflammatory cytokines including CCl12, CCL17, CCL24, and Eotaxin in the mouse lung digests as compared to air control. Interestingly, the fold changes in the PG:VG+Nic group were contrasting to those observed by PG:VG alone and flavored e-cig aerosol exposed mouse lungs, but none of these changes were highly significant.

To identify genes that were commonly altered upon exposure, we generated a list of common genes that were significantly dysregulated in exposure categories (fruit, menthol and tobacco). We identified 9 such target genes-Ne*url3*, *Egfem1*, *Stap1*, *Tfec*, *Mitf*, *Cirbp*, *Hist1h1c*, *Gmds,* and *Htr2c-* that were dysregulated in the myeloid cluster from lungs exposed to differently flavored e-cig aerosol, but not PG:VG. We observed significant upregulation of *Neurl3*, *Stap1*, *Cirbp,* and *Hist1h1c* and downregulation of *Tfec*, *Mitf*, *Gmds,* and *Htr2c* in the myeloid cluster of mice exposed to differently flavored e-cig aerosols. Upon analyzing the lymphoid cluster for commonly dysregulated genes, we identified – *Klra8* (Killer cell lectin-like receptor 8) and *Nfia* (nuclear factor I)-that were significantly upregulated in the exposure groups as compared to air-controls **(Figure S5A)**. Klra8 is a natural killer cell associated gene, and its upregulation is generally associated with viral infection associated host immune responses within the mouse lungs (32–34). *Nfia* is a transcriptional activator responsible for regulating Oxphos-mediated mitochondrial responses and proinflammatory pathways (35, 36).

Overall, we identified a total of 29 commonly dysregulated gene targets that were identified from five major cell clusters and performed gene enrichment analyses on the identified targets to identify the top hits **(Tables 2-3)**. Terms like ‘negative regulation of immune system’ (*Hmgb3*/*Gpam*/*Scgb1a1*/*Stap1*/*Ldlr*), ‘positive regulation of lipid biosynthetic pathway’ (*Htr2c*/*Gpam*/*Ldlr*), and ‘receptor recycling’ (*Ldlr*/*Ramp3*) were amongst the top hits in our observations **(Figure S5B)**.

**Table 2:**
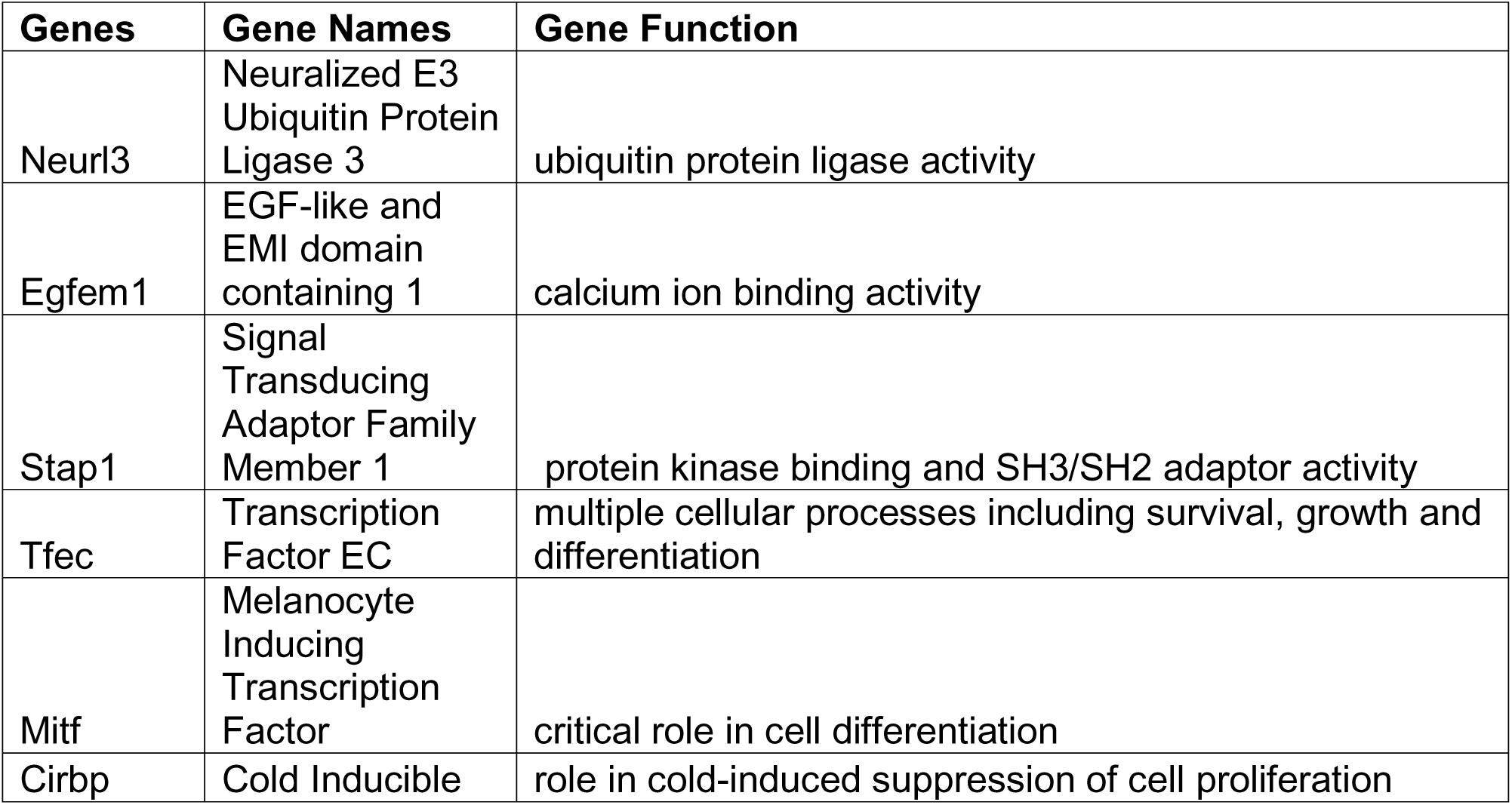
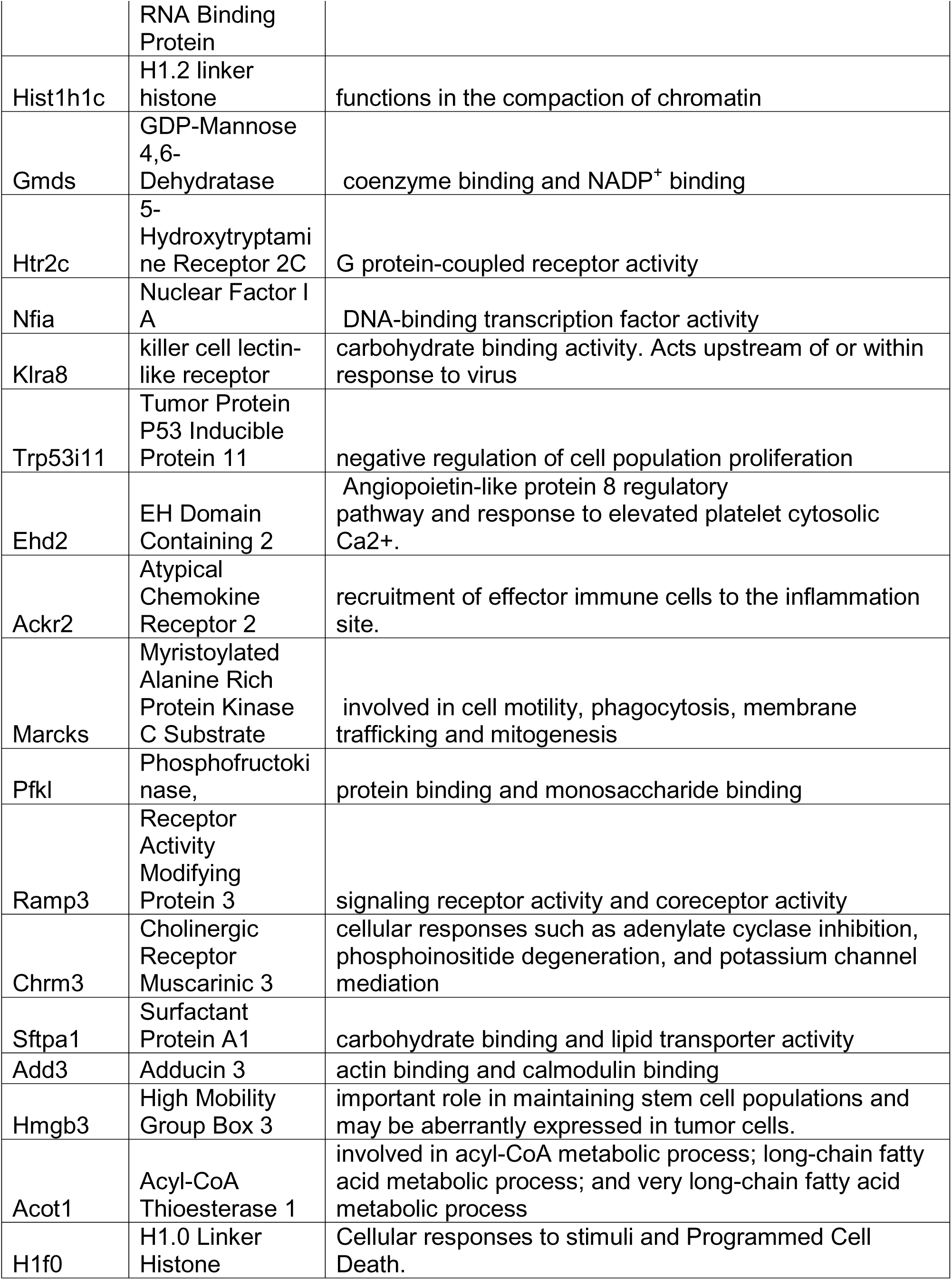
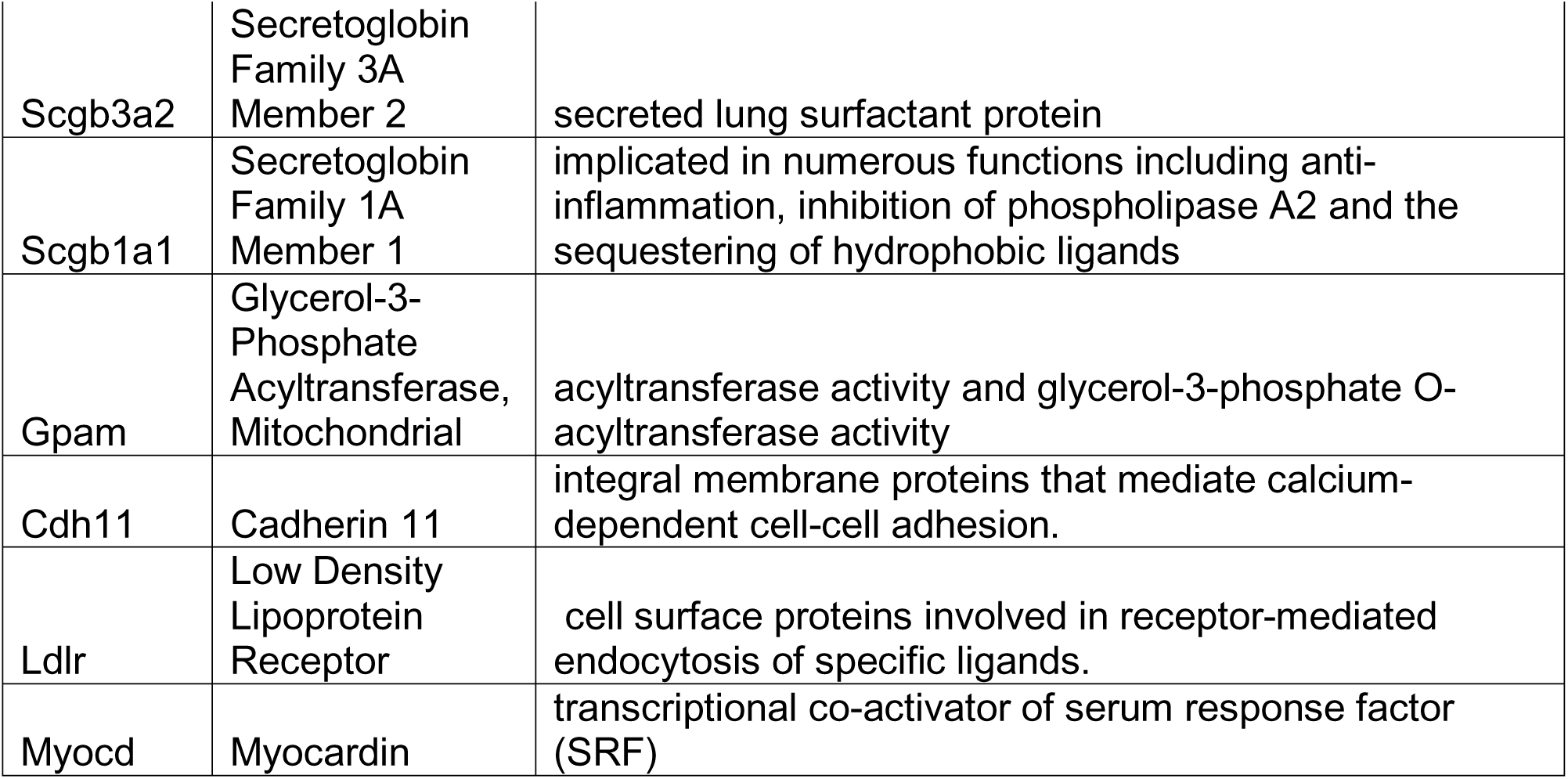
List of top dysregulated genes on exposure to differently flavored (fruit, menthol and tobacco) e-cig aerosol in C57BL/6J mouse lungs.

**Table 3:**
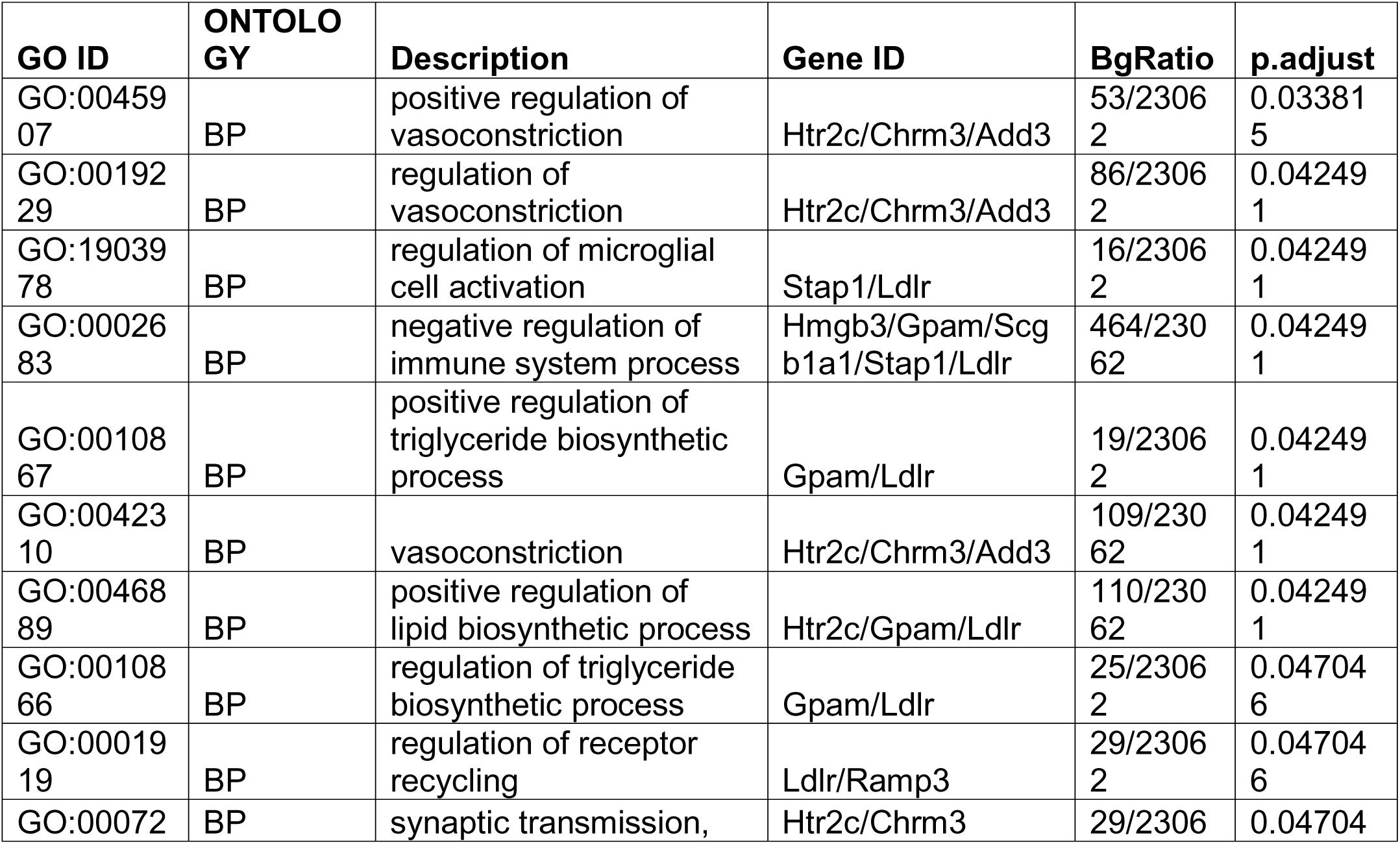
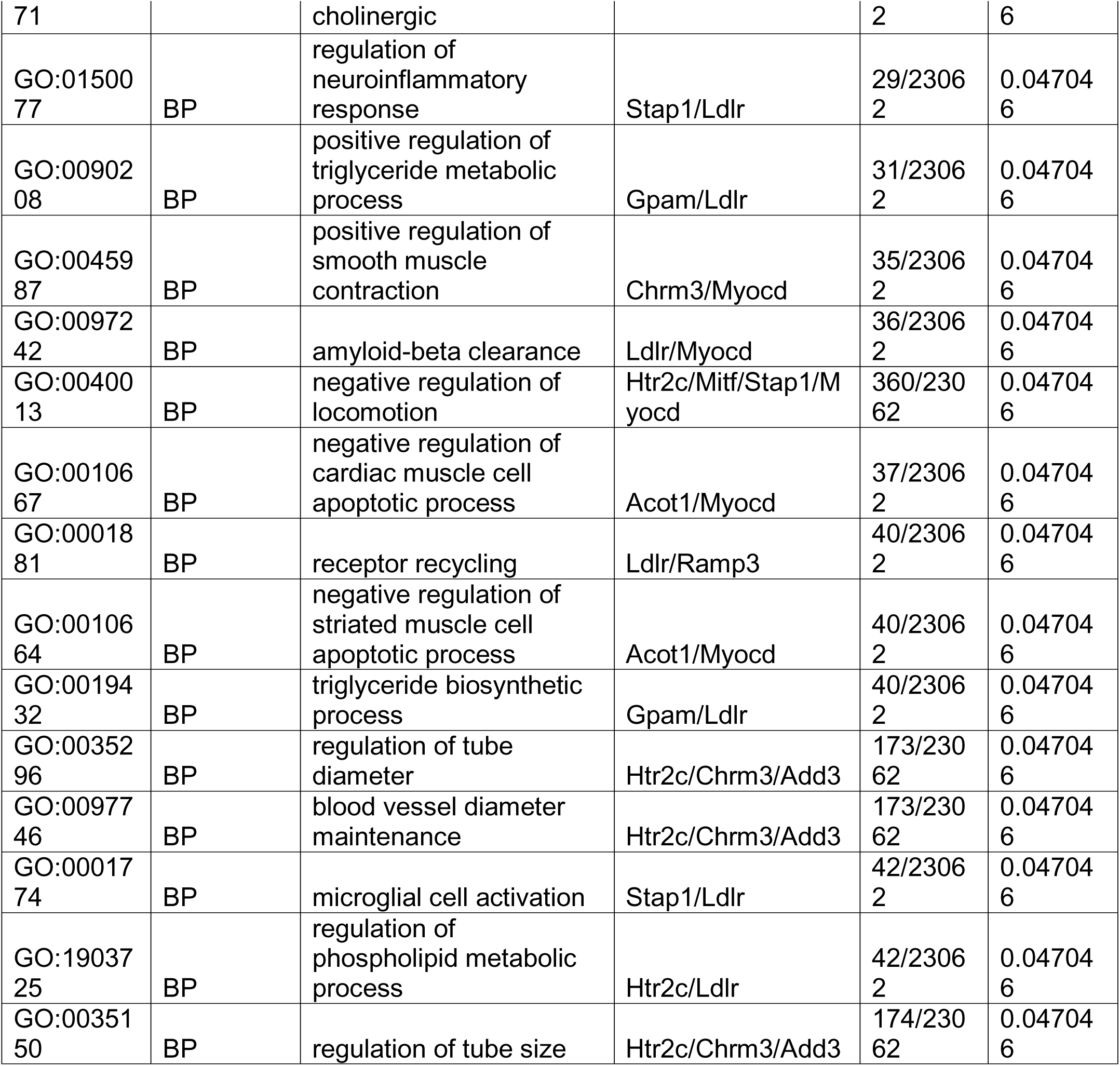
Gene Ontology results showing the top hits from the commonly dysregulated genes in all clusters on exposure to e-cig aerosols.

Of note, the data presented in this study is a sub-part of a larger study. In addition to the groups mentioned in this manuscript, we also had two additional groups of TDN and Tobacco-Free Nicotine (TFN). Though further objectives and experimentations performed in both these studies were varied, common air and PG:VG samples were used for analyses of cytokine/chemokine and cell count data as described in our previous publication(24).

## Discussion

E-cigs and associated products have constantly been under scrutiny by the US Food and Drug Administration (FDA) due to public health concerns. In February 2020, FDA placed a regulation on all cartridge-based flavored e-cigs except for menthol and tobacco to reduce the use of e-cigs amongst adolescents and young adults. But it left a loophole for the sale of flavored (including menthol) disposable and open system e-cigs (1, 37). Importantly, most e-cig related bans in the US happened at the state level, thus allowing differential levels of restrictions imposed on the premarket tobacco applications (PMTAs) and sales which defeats the purpose of limiting their accessibility to the general public (38, 39). In fact, the use of nicotine containing e-cigs amongst youth and associated policy restrictions have recently been found to be linked to unintended increase in traditional cigarette use (40). Each year new products are introduced in the market with newer device designs and properties to lure the users (adults between the ages of 18-24 years) which makes it crucial to continue with the assessments of toxicity and health effects of e-cigs in an unbiased manner (41).

Numerous studies indicate increased oxidative stress, DNA damage, and loss of neutrophil function due to exposure to e-cig aerosols *in vitro* and *in vivo* (7, 24, 42–45). However, we do not have much knowledge about the cell populations and biological signaling mechanisms that are most affected upon exposure to differently flavored e-cig aerosols at a single-cell level. To bridge this gap in knowledge, we studied the transcriptional changes in the inflammatory responses due to acute (1-hr nose-only exposure including 120 puffs for 5-consecutive days) exposure to fruit-, menthol– and tobacco-flavored e-cig aerosols using single-cell technology. Short-term (2 hr/day for 3 or 5 consecutive days) e-cigarette exposure studies conducted by others and our group have shown increase in the pro-inflammatory cytokines and oxidative stress markers in both pulmonary and vascular tissues (25, 46, 47). Since nose-only exposures are more direct than whole-body, we reduced the daily exposures of mice to one hour for 5 consecutive days. This dose and duration was found to show sex-dependent changes in MMP-2 and –9 activity and expression (24). Prior literature formed the basis of the current study to assess the effect of acute exposure to differently flavored e-cig aerosols on the lung cell population using a nose-only exposure system.

Reports indicate that the release of metal ions due to the burning of metal coil is a major source of variation during e-cig exposures (48–50). A 2021 study reported the presence of 21 elements in the pod atomizers from different manufacturers identifying a high abundance of 11 elements including nickel, iron, zinc, and sodium amongst others (51). Previous studies have also shown the presence of similar elements in the e-cig aerosols which could have a possible adverse health effect on the vapers (19, 52). More importantly, our study points towards a much important issue, which pertains to the product design of the e-cig vapes. We believe that the aerosol composition varies based on the type of atomizer, coil resistance, coil composition, and chemical reactivity of the e-liquid being used. While much work is done on the chemical composition of the flavors and e-liquids, the other aspects of device design remain understudied and must be an area of research in the future. In fact, a recent publication by our group emphasizes this aspect of product design by studying the exposure profile from low and high-resistance coil in an e-cig (21). Another factor that may limit the interpretation of our results pertains to the correlation between the leached metal and the observed transcriptional changes. Since our study provides proof of day-to-day variation in the leaching of metal ions from the same liquid using the same atomizer in the emitted aerosols, it could be possible to develop a statistical model correlating the differential metal exposure to the gene expression changes to get a good assessment of the dose of each toxicant deposited on the lung surface, which has not been done in this study. This raises a pertinent question about what parameters should be controlled to design a comparative study between different flavors of e-cigs in the market per the standards of particle distribution and characteristics. Flavor-dependent variations in daily leaching of metals from an e-cig coil may result in variable deposition of toxicants onto the epithelial lining of both mouth and lungs in frequent vapers. Thus, future studies need to consider conditions like flavor, wattage, coil resistance, coil composition and atomizer life when designing *in vitro* and *in vivo* studies to deduce the acute and chronic toxicities of vaping in human.

E-cig vaping has been known to affect the innate and adaptive immune responses among vapers (44, 53–55), but flavor-specific effects on immune function are not fully explored. While we expected to see flavor-dependent changes in our experiment, we did not anticipate observing interesting sex-dependent variation in the lung tissues using flow cytometry. In this respect, recent studies show concurring evidence suggesting sex-specific changes in lung inflammation, mitochondrial damage, gene expression, and even DNA methylation in mice exposed to e-cig aerosols (56, 57).

It is important to highlight here that while the changes in macrophage and neutrophil frequencies ascertained by scRNA seq corroborate with that observed using flow cytometry; similar correlation was not observed for the CD4^+^ and CD8^+^ T cell data. The possible explanation for such a discrepancy could be high gene dropout rates in scRNA seq (58), different analytical resolution for the two techniques (59) and pooling of samples in our single cell workflow. Also, while we were able to identify some interesting changes in the eosinophil population within the lungs upon exposure to e-cigs using our flow cytometry data, we could not identify a cluster for eosinophils in our scRNA seq dataset. This could be due to the loss of this cell type during the sample preparation during scRNA seq capture or filtering out as empty droplets during data analyses. Theoretically, eosinophils are known to have lower RNA content and express fewer number of genes, which may result in them being identified as empty droplets and removed when running Cell Ranger (60). Despite the lack of data from scRNA seq, our findings are of relevance as an increased neutrophilic, but a decreased eosinophilic response is characteristic of severe inflammation, infection and asthma (61–63). Further probing into the role of these two cell types in shaping the immune landscape within the lung upon e-cig aerosol exposure is paramount in understanding the specific toxicities and impact of each e-cig flavor on human health and well-being.

One of the most interesting discoveries from our single-cell analyses was the identification of a cluster of immature neutrophils without Ly6G surface marker. While we observed an increase in the cell percentages of these cells in our treatment groups, little to no change in the gene expression was noted (data not shown) which could be indicative of impaired function of mature neutrophil population, a fate being reported by various previous studies pertaining to e-cig exposures (13, 53). In fact, our study identified (a) a moderate increase in the neutrophil count in menthol and tobacco-flavored e-cig aerosol treated mouse lungs through scRNA seq analyses, (b) corroboration of the findings from scRNA seq using flow cytometry with more pronounced increase in Ly6G+ neutrophils for male menthol-flavored e-cig aerosol exposed mice and (c) identification of decreased co-localization for Ly6G and S100A8 in the lungs of tobacco-flavored e-cig aerosols as compared to air control through co-immunostaining. Ly6G is an important marker of neutrophil recruitment and maturation in mammalian cells and has been reported in relation to various bacterial and parasitic infections in previous studies (64, 65). Of note, this cluster did not express SiglecF and had high expression of other neutrophil markers including S100A8/9, Lcn2 and Il1b, thus negating the possibility of eosinophils being misrepresented as ‘Ly6G-neutrophils’ in our analyses.

S100A8 act as damage associated molecular patterns (DAMPs) and are responsible for neutrophil activation and neutrophil extracellular trap (NET) formation (66, 67). While scRNA seq results in our study identified expression of S100A8/A9 genes in both neutrophil clusters –Ly6G+ and Ly6G-, the expression of Ly6G was totally absent for clusters identified as Ly6G-. Co-immunoprecipitation results also showed expression of both S100A8 and Ly6G markers within the lungs of treated and control lungs, but the co-localization of the two markers was more prominent in the control lungs, thus pointing towards a possible shift in the neutrophil function and activity upon exposure to e-cig aerosols. We are not the first to report the Ly6G deficiency amongst neutrophils.

Previous work by Deniset et al (2017) and Kleinholz et al (2021) have highlighted the importance of Ly6G deficiency in relation to infection (64, 65). Deniset et al in their 2017 study reported the presence of two populations of neutrophils in the splenic tissue of *Streptococcus pneumoniae* infected mice, based on the Ly6G expression– Ly6G^high^ and Ly6G^intermediate^. They found that while the former corresponds to mature neutrophils with ability of tissue migration and bacterial clearance, the latter constitutes the immature, immobile pool of neutrophils responsible for proliferation and replenishment of the mature pool of neutrophils (64). The later study by Kleinholz et al (2021) demonstrated decreased uptake of *Leishmania major* in Ly6G deficient mice thus leading to delayed recruitment to and pathogen capture by neutrophils at the site of infection. Taken together these studies, provide evidence for a protective/compensatory role of the loss of Ly6G in the neutrophil population (65). This in addition to the recent findings by Jasper et al (2025) where neutrophils from healthy volunteers demonstrated a reduction in neutrophil chemotaxis, phagocytic function, and NET formation upon exposure to e-cig aerosols (44), point towards a probable shift in the neutrophil dynamics upon exposure to e-cig aerosols. It is important to mention here that though our results from co-immunostaining using Ly6G and S100A8 pointed towards a shift in innate immune responses upon exposure to e-cig aerosols, it must not be confused with them being only expressed by neutrophil population. S100A8 is expressed in myeloid population including neutrophils, monocytes and macrophages (68, 69) and a subgroup of eosinophils are also known to express Ly6G (70). Thus, an in-depth characterization of these identified population is important to understand the cellular and molecular responses towards e-cig aerosol exposure *in vivo*.

The myeloid and lymphoid limbs of immunity are interconnected (71, 72). In our study, we find an incidental shift in the neutrophil dynamics in menthol and tobacco flavored e-cig exposed mouse lungs. In contrast, we report an increase in the T-cell responses in the form of increased CD8+ T cells from both scRNA seq and flow cytometric analyses in male mice. In fact, increased expression of genes including *Malt1*, *Serpinb9b,* and *Sema4c* are indicative of enhanced T-cell mediated immune response in the lungs of mice exposed to fruit (mango) flavored e-cig aerosols (73–75). Contrary to this, exposure to menthol-flavored e-cig aerosols had a much milder effect on the lymphoid population within the lungs of C57BL/6J mice. We found evidence for increased lymphoid cell proliferation due to activation of cyclin-dependent protein kinase signaling mediated via expression of genes including *Cdk8* and *Camk1d* in these cells (29, 76). Exposure to tobacco-flavored e-cig aerosol provided evidence for decreased chaperone-mediated protein folding, due to the downregulation of Chaperonin Containing TCP-1 (CCT) family of proteins. Chaperonin Containing TCP-1 proteins are important to regulate the production of native actin, tubulin, and other proteins crucial for cell cycle progression and cytoskeletal organization (77). This is in conjunction with the upregulation of *Klra4* and *Klra8* that is indicative of increased protein misfolding and cytotoxic responses in the lymphoid cells of tobacco-exposed e-cig aerosols (78, 79).

Overall, we provide evidence of altered innate immune responses due to variable neutrophilic-eosinophilic function and increased T-cell proliferation and cytotoxicity in a flavor-dependent manner upon exposure to e-cig aerosols in this study. An increase in the levels of CCL17, CCL20, CCL22, IL2 and Eotaxin in the lung digests from tobacco-exposed mouse lungs further support this deduction as these cytokines/chemokines are associated with T-cell mediated immune responses (80–87). Importantly CXCL16 attracts T-cells and natural killer cells to activate cell death. It is involved in LPS-mediated acute lung injury (ALI), an outcome which has also been linked with e-cig exposures in human (88, 89).

Further, we compiled a list of commonly dysregulated genes in a flavor independent manner and identified 29 gene targets. Signal-transducing adaptor protein-2 (*Stap1)* which is commonly upregulated upon exposure to e-cig aerosols is known to regulate T-cell activation and airway inflammation which is in agreement with the overall outcome of our findings (90). Another gene that was found to be consistently dysregulated in many cell types was Cold-inducible RNA binding protein (*Cirbp*). *Cirbp* is a stress response protein linked with stressors like hypoxia. Its upregulation upon e-cig exposure supports that vaping induces oxidative stress and can have adverse implications on the exposed cell types (91). Importantly, this gene is involved in DNA repair mechanism thus making it crucial for cell survival pathways (92). 5-hydroxytryptamine receptor 2C (*Htr2c*) and *Klra8* are other genes in this category of commonly dysregulated genes that are associated with enhancing inflammation and cell death (78, 79, 93). Overall, we provide a cell-specific resource of immune responses upon exposure to differently flavored e-cig aerosols.

Considering that scRNA technology has not been commonly used for e-cig research, ours is one of the first studies employing this technique to identify possible changes in the cellular composition and gene expressions. Importantly we use the nose-only exposure system for our experiment to avoid exposure through other routes. However, despite the novel approach and state-of-the-art exposure system, we had a few limitations. First, we used a small sample size to identify the changes in the mouse lungs upon exposure to e-cig aerosols at a single-cell level. Due to the expensive nature of single cell sequencing technology and limited information in literature, we chose to design this experiment with small sizes of experimental and validation cohort. But, based on the encouraging findings from this study, future studies could be designed with a larger sample size, longer durations of exposure and more targeted approach to identify the acute and chronic effects of vaping *in vivo*. Second, we could not expand upon the sex-dependent changes observed through our work upon exposure. This was because such an effect was not anticipated when we conceived the idea of a short-term exposure in mice. However, considering the evidence from the current study, future experimental designs in our lab are considering sex as a crucial confounder for studying the effects of e-cig exposure in translational contexts. Third, the inclusion of PG:VG + Nic group was streamlined in this study, but in future work inclusion of this group for scRNA seq analyses to delineate the effects of nicotine alone on gene transcription is necessary. Fourth, we did not anticipate changes in the metal release on consecutive days of exposure at the start of our study. Later, our data pointed towards the importance of device design in e-cig exposures. Future studies need to identify the factors that may affect the daily composition of e-cig aerosols and devise a method of better monitoring these possible confounders. However, in this regard our experiment does mimic the real-life scenario, as such variations due to prolonged storage of e-liquid and differences arising due to vape design must be common amongst human vapers.

In conclusion, we identified cell-specific changes in the gene expressions upon exposure to e-cig aerosols using single cell technology. We identified a set of top 29 dysregulated genes that could be studied as markers of toxicity/immune dysfunction in e-cig research. Future work with larger sample sizes and sex-distribution is warranted to understand the health impacts of long-term use of these novel products in humans.

## Methods

### Animals Ethics Statement

All experiments were conducted per the guidelines set by the University Committee on Animal Resources at the University of Rochester Medical Center (URMC). Care was taken to implement an unbiased and robust approach during the experimental design and conduction of each experiment to ensure data reproducibility per the National Institutes of Health standards.

### Animals

We ordered 5-week-old pups of male and female C57BL/6J mice from Jackson Laboratory to conduct this experiment. Prior to the start of the experiment, mice were housed at the URMC Vivarium for acclimatization. Thereafter the animals were moved to the mouse Inhalation Facility at URMC for training and exposures.

One week prior to the start of the exposures, mice underwent a five-day nose-only training to adapt themselves to the mesh restraints of the exposure tower. The mouse restraint durations were increased gradually to minimize the animal’s stress and discomfort. Of note, the mouse sacrifice was performed within 8-12 weeks’ age for each mouse group to ensure that the mouse age corresponds to the age of adolescents (12-17 years) in humans (94, 95). Age and sex-matched animals (n = 2/sex/group) used to perform single cell RNA sequencing (scRNA seq) were considered as the ‘experimental cohort’; whereas another group of age and sex-matched (n = 3/sex/group) mice exposed to air and flavored e-cig aerosol served as ‘validation cohort’ for this study.

### E-cigarette Device and E-liquid

We utilized an eVic-VTC mini and CUBIS pro atomizer (SCIREQ, Montreal, Canada) with a BF SS316 1.0-ohm coil from Joyetech (Joyetech, Shenzhen, China) for vaping and the inExpose nose-only inhalation system from SCIREQ (SCIREQ, Montreal, Canada) for mouse exposures. Both air and PG:VG exposed mice groups were considered as controls for this experiment. We used commercially available propylene glycol (PG; EC Blend) and vegetable glycerin (VG; EC Blend) in equal volumes to prepare a 50:50 solution of PG:VG. For flavored product exposures, mice were exposed to three different e-liquids – a menthol flavor “Menthol-Mint”, a fruit flavor “Mango” and a tobacco flavor “Cuban Blend”. Of note, all the e-liquids were commercially manufactured with 50mg/mL of tobacco derived nicotine (TDN). So, all treatments have nicotine in addition to the flavoring mentioned respectively. Additionally, we used a mixture of PG:VG with 50mg/mL of TDN as a control for limited experiments to study the effect of nicotine alone in our treatment. This group is labeled PG:VG+Nic for the rest of the manuscript.

### E-cigarette Exposure

Scireq Flexiware software with the InExpose Inhalation system was used for controlling the Joyetech eVic-VTC mini device to perform nose-only mouse exposures. For this exposure, we utilized a puffing profile that mimicked the puffing topography of e-cig users in two puffs per minute with a puff volume of 51 mL, puff duration of 3 seconds, and an inter puff interval of 27 seconds with a 2 L/min bias flow between puffs (96). **Figures 1A** & **S1A** depict the experimental design and exposure system employed for this study.

Age-matched male and female (n=5 per sex) mice were used for each group, namely, air, PG:VG, Fruit, Menthol, and Tobacco. To ensure rigor and reproducibility in our work, we have used age– and sex-matched control and treated mice in this study. Confounders like environment and stress were minimized by housing all the cages in environmentally controlled conditions and training all the mice (both control and treated) in nose-only chambers. Each group of mice was exposed to the above-mentioned puffing profile for one hour each day (120 puffs) for a total of five consecutive days. Additionally, a group of mice (n=3/sex) was exposed to PG:VG + Nic for the same duration using similar exposure profile to serve as control to assess the effect of nicotine on the observed changes using selected experiments. Air-exposed mice were exposed to the same puffing profile for a total of five consecutive days to ambient air. We recorded the temperature, humidity, and CO levels of the aerosols generated at the start, mid, and end of the exposure on each day using the Q-Trak Indoor Air Quality Monitor 7575 (TSI, Shoreview, MN). Total Particulate Matter (TPM) sampling was done from the exhaust tubing of the set-up at the 30 min mark of the exposure and at the inlet connected to the nose-only tower (shown in **Figure S1A**) immediately after the culmination of the exposure. Gravimetric measurements for TPM were also conducted to confirm relative dosage to each mouse group daily.

### Preparation of single-cell suspension

The animals (at 8-10 weeks’ age) were sacrificed immediately after the final exposure. Vascular lung perfusion was performed using 3 mL of saline before harvesting the lung lobes for preparation of single-cell suspension. It is important to mention here that of the 5 lung samples/sex/group; 2 sex/group were used for histological assessments and scRNA analyses. Here, the left lung lobe was inflated using low-melting agarose and used for histology, while the rest of the uninflated lung lobes were used for preparing the single-cell suspension. We pooled the lung lobes for each sex per group for preparation of the single-cell suspension as depicted in **Figure 1A**. The lung lobes were weighed and digested using Liberase method as described earlier (97). Briefly, lung lobes were weighed and digested using Liberase (Cat# 5401127001; Roche, Basel, Switzerland) enzymatic cocktail with 1% DNase. The tubes were then transferred to the gentleMACS dissociator (Miltenyi, Gaithersburg, MD) and the manufacturer’s protocol for mouse lung digestion was run. The sample tubes were next incubated at 37^0^C for 30 minutes with constant rotation after which the suspension was strained through 70-micron MACS Smart Strainer. Thereafter, the suspension was centrifuged at 500g for 10 minutes at 4^0^C, the supernatant was discarded, and 0.5 mL of RBC lysis buffer was added to the cell pellet to digest RBCs. The suspension was left on ice for 5 min in RBC lysis buffer and then 4 mL of ice-cold PBS with 10% FBS was added to stop the lysis. The suspension was again centrifuged at 500g for 10 minutes at 4^°^C. The cell pellet was suspended in 1 mL PBS with 10% FBS, and cell number and viability were checked using AO/PI staining on a Nexcelom Cellometer Auto2000.

### Library Preparation and Single-Cell Sequencing

The prepared single-cell suspension was sent to the Genomics Research Center (GRC) at URMC for library preparation and single-cell sequencing. Library preparation was performed from control and treatment groups using the 10X single cell sequencing pipeline by 10X Genomics and 10,000 cells were captured per sample using the Chromium platform. The prepared library was sequenced on NovaSeq 6000 (Illumina, San Diego, CA) at a mean sequence depth of 30,000 reads per cell. Read alignment was performed to GRCm38 Sequence.

### Data Analyses

We used the standard Seurat v4.3 analyses pipeline to analyze our data (98). In brief the low-quality cells and potential doublets were excluded from the dataset to create the analyses dataset. The residual features due to the presence of RBCs were corrected before integration. “scTransform” function was used for integration of all the datasets after which the standard Seurat pipeline was used for data normalization of integrated data. “FindVariableGenes” gene function was used to identify the variable genes for dimensionality reduction using PCA function. UMAP was used for dimensionality reduction and clustering of cells.

To identify the unique features and cell clusters within each cell subtype, we used the sub-setting feature within Seurat. After identifying the 5 major cell populations (epithelial, endothelial, stromal, myeloid and lymphoid) in our data sets, each of these cell types were sub-clustered using “subset” function, normalized and re-clustered. Cell annotation for each of the subsets was performed with the help of Tabula Muris database (99, 100). However, some clusters were annotated manually with the help of a literature search as discussed in the results section.

DESeq2 (V.1.42.1) was used to perform pseudobulk analyses to identify differentially expressed genes within each group. Here, genes showing a fold change > 0.5 & < –0.5 along with a p_adj_ value < 0.05 were considered significantly dysregulated and plotted as heatmap using GraphPad The ClusterProfiler R package (V. 4.10.1) (101) was employed to perform gene enrichment analyses of the differentially expressed genes (fold change> 0.5 & < –0.5).

### Cytokine/chemokine assessment

We used multiplex assay to determine the levels of cytokine/chemokine in the lung homogenates from control and e-cig aerosol exposed mouse lungs using commercially available Bio-Plex Pro Mouse Chemokine Assay (Cat#12009159, Bio-RAD, Hercules, CA) per the manufacturer’s instructions. Approximately 40 mg of mouse lung lobes were homogenized in 300uL of 1X RIPA buffer with 0.1% protease and phosphatase inhibitor. The lung homogenate was stored on ice for 30 min. Following incubation, the homogenate was centrifuged at 15000 rpm for 15 min at 4 degrees Celsius. The supernatant was collected and used for performing the multianalyte assay for determination of cytokine/chemokine levels using Luminex FlexMap3D system. A heatmap after normalization of the measured cytokine/chemokine to the protein amount loaded was plotted.

### Lung Histology

The left lung lobe of mice used for scRNA seq were inflated with 1% low melting agarose and fixed with 4% neutral buffered PFA. Fixed lungs were dehydrated, before being paraffin-embedded and sectioned (5μm). Hematoxylin and eosin (H&E) staining was performed by the Histology, Biochemistry, and Molecular Imaging Core at URMC. The H&E stain was observed at 10X magnification using Nikon Elipse-Ni fluorescence microscope. Ten to fifteen random images were captured per sample.

### Flow cytometry

Flow cytometry was performed on the cells collected from lung homogenates from air and flavored e-cig aerosol exposed mouse lungs. For analyses of immune cell population in the lung, the lung lobes were digested as described earlier (97). The single-cell suspension thus prepared was used to run flow cytometry using the BD LSRFortessa cell analyzer. Cells were blocked with CD16/32 (Tonbo biosciences 70-0161-u500, 1:10) to prevent nonspecific binding and stained with a master mix of Siglec F (BD OptiBuild Cat#740280, 1:200), CD11b (Biolegend Cat #101243, 1:200), Ly6G (BD Horizon Cat# 562700, 1:200), CD45 (Biolegend Cat#103126, 1:200), CD11c (Biolegend Cat #117318, 1:200), CD4 (Biolegend Cat#116012, 1:200), and CD8 (eBiosciences Cat#17-0081-82, 1:200). 7AAD (eBiosciences Cat#00-6993-50, 1:10) was used as the nucleic acid dye to detect live and dead cells. The gating strategy used for this assay has been depicted in **Figure S6**.

### Metal analyses

To understand the levels of metals released during subsequent days of exposure, we performed Inductively coupled plasma mass spectrometry (ICP-MS) on the e-cig aerosol condensates collected from each day of exposure using a Perkin Elmer ICP-MS model 2000C. The samples were run using a Total Quant KED protocol with 4mL/min Helium flow and externally calibrated using a blank and a 100ppb standard for the 51 elements. The samples were submitted to the Element Analyses facility at URMC, and levels of metals thus detected were plotted.

### Ly6G /S100A8 Double staining

To determine the various populations of neutrophils in exposed and control groups, FFPE tissue sections from air and tobacco-flavored e-cig aerosol exposed lungs were stained with Ly6G and S100A8. In brief, 2-3 tissue sections per sample were deparaffinized using serial incubation in xylene followed by graded alcohol. Slides were incubated in 1X Citrate Buffer (Cat# S1699, Agilent, Santa Clara, CA) for 10 min at 95°C for antigen retrieval which was followed by incubation at room temperature for 30 min. The slides were next washed with water and permeabilized using a permeabilization buffer (0.1% Triton-X in 1X TBST) for 10 min. Next, the slides were again washed with 1X TBST and blocked using Blocking buffer (5% goat serum in 1X TBST) for 30 min at room temperature. The blocked slides were incubated overnight at 4°C with Ly6G (Cat# 16-9668-85, Invitrogen, dilution: 1:100) and S100A8 (Cat# 26992-1-AP, Proteintech, dilution: 1:200). Next day, the slides were washed with 1X TBST and incubated for 2 hrs. at room temperature with goat anti-rabbit Alexa Fluor 594 (Cat # A11012, Invitrogen) and donkey anti-mouse Alexa Fluor 488 (Cat # A21202, Invitrogen) secondary antibody at 1:1000 dilution. Thereafter the slides were washed and mounted with ProLong Diamond Antifade Mountant with DAPI (Cat# P36962, Invitrogen, Waltham, MA). 6-10 images were captured using Cytation 5 Imaging software (Thermo Fisher, Waltham, MA) at 10X magnification. The ImageJ deconvolution was used for quantifying the fluorescence in green (Ly6G+) and red (S100A8+) channels relative to DAPI (blue channel) and the relative fluorescence was plotted.

### Statistical significance

We used GraphPad Prism 10.5.0 for all statistical calculations. All the data plotted in this paper are expressed as mean ± SEM. Pairwise comparisons were done using unpaired *t* test while one-way analysis of variance (ANOVA) with ad-hoc Tukey’s test was employed for multi-group comparisons. To identify sex-based variations in our treatment groups, Tukey post hoc two-way ANOVA was employed.

### Data Availability

The datasets generated during and/or analyzed during the current study are deposited on NCBI Gene Expression Omnibus. Specifically, mouse scRNAseq is available under accession code “GSE263903” (https://www.ncbi.nlm.nih.gov/geo/query/acc.cgi?acc=GSE263903). The data will be publicly available upon publication or on April 11, 2026, whichever is earlier. Reviewer’s token for access is ‘wtofqmayjdeltcn’. All other relevant data supporting the key findings of this study are available within the article and its Supplementary Information files or from the corresponding author upon reasonable request. Source data are provided with this paper.

### Code Availability

Data collection was performed with mkfastq pipeline in Cell Ranger’s (v7.0.1). Cell Ranger (v7.0.1) was used for cell and gene counting using the default settings. Single-cell analysis was performed using the Seurat R package (v4.3.0) using the recommended workflow.

### Author’s contributions

GK, TL and AT planned and conducted the experiments; TL analyzed the lab-based assays; GK and AT analyzed the scRNA data; GK wrote and edited the manuscript; TL, AT and IR reviewed the data and edited the manuscript; IR conceived and procured funding for the project.

## Acknowledgements & Funding Sources

We would like to thank the Genomics Research Core, the Elemental Analyses Facility, and the Histology, Biochemistry, and Molecular Imaging Core at URMC for assisting us in the scRNA seq, metal analyses in aerosols, and lung sectioning and histology respectively. We would also like to acknowledge Chengru Jiang for helping with histology image acquisition for this manuscript. This work was supported by WNY Center for Research on Flavored Tobacco Products (CRoFT) # U54CA228110 and Toxicology Training Program T32 ES007026.

## Conflict of Interest

The authors have no financial disclosure or conflict of interest with the findings presented in this research article.

## Notes

### Competing Interest Statement

The authors have declared no competing interest.

### Summary of Updates

Second Revision to Responses based on reviewers comments.

## References

1. Ma S, Qiu Z, Yang Q, Bridges JFP, Chen J, Shang C. Expanding the E-Liquid Flavor Wheel: Classification of Emerging E-Liquid Flavors in Online Vape Shops. Int J Environ Res Public Health. 2022;19(21).

2. Wang TW, Neff LJ, Park-Lee E, Ren C, Cullen KA, King BA. E-cigarette Use Among Middle and High School Students – United States, 2020. MMWR Morb Mortal Wkly Rep. 2020;69(37):1310–2.

3. Wu D, O’Shea DF. Potential for release of pulmonary toxic ketene from vaping pyrolysis of vitamin E acetate. Proc Natl Acad Sci U S A. 2020;117(12):6349–55.

4. Goniewicz ML, Kuma T, Gawron M, Knysak J, Kosmider L. Nicotine levels in electronic cigarettes. Nicotine Tob Res. 2013;15(1):158–66.

5. Lee YJ, Na CJ, Botao L, Kim KH, Son YS. Quantitative insights into major constituents contained in or released by electronic cigarettes: Propylene glycol, vegetable glycerin, and nicotine. Sci Total Environ. 2020;703:134567.

6. Wang Q, Sundar IK, Blum JL, Ratner JR, Lucas JH, Chuang TD, et al. Prenatal Exposure to Electronic-Cigarette Aerosols Leads to Sex-Dependent Pulmonary Extracellular-Matrix Remodeling and Myogenesis in Offspring Mice. Am J Respir Cell Mol Biol. 2020;63(6):794–805.

7. Muthumalage T, Lamb T, Friedman MR, Rahman I. E-cigarette flavored pods induce inflammation, epithelial barrier dysfunction, and DNA damage in lung epithelial cells and monocytes. Sci Rep. 2019;9(1):19035.

8. Lee HW, Park SH, Weng MW, Wang HT, Huang WC, Lepor H, et al. E-cigarette smoke damages DNA and reduces repair activity in mouse lung, heart, and bladder as well as in human lung and bladder cells. Proc Natl Acad Sci U S A. 2018;115(7):E1560–e9.

9. White AV, Wambui DW, Pokhrel LR. Risk assessment of inhaled diacetyl from electronic cigarette use among teens and adults. Sci Total Environ. 2021;772:145486.

10. Martin EM, Clapp PW, Rebuli ME, Pawlak EA, Glista-Baker E, Benowitz NL, et al. E-cigarette use results in suppression of immune and inflammatory-response genes in nasal epithelial cells similar to cigarette smoke. Am J Physiol Lung Cell Mol Physiol. 2016;311(1):L135–44.

11. Sussan TE, Gajghate S, Thimmulappa RK, Ma J, Kim JH, Sudini K, et al. Exposure to electronic cigarettes impairs pulmonary anti-bacterial and anti-viral defenses in a mouse model. PLoS One. 2015;10(2):e0116861.

12. Masso-Silva JA, Moshensky A, Shin J, Olay J, Nilaad S, Advani I, et al. Chronic E-Cigarette Aerosol Inhalation Alters the Immune State of the Lungs and Increases ACE2 Expression, Raising Concern for Altered Response and Susceptibility to SARS-CoV-2. Front Physiol. 2021;12:649604.

13. Madison MC, Landers CT, Gu BH, Chang CY, Tung HY, You R, et al. Electronic cigarettes disrupt lung lipid homeostasis and innate immunity independent of nicotine. J Clin Invest. 2019;129(10):4290–304.

14. Cao Y, Wu D, Ma Y, Ma X, Wang S, Li F, et al. Toxicity of electronic cigarettes: A general review of the origins, health hazards, and toxicity mechanisms. Sci Total Environ. 2021;772:145475.

15. Jovic D, Liang X, Zeng H, Lin L, Xu F, Luo Y. Single-cell RNA sequencing technologies and applications: A brief overview. Clin Transl Med. 2022;12(3):e694.

16. Inayatullah M, Dwivedi AK, Tiwari VK. Advances in single-cell omics: Transformative applications in basic and clinical research. Current Opinion in Cell Biology. 2025;95:102548.

17. Ke M, Elshenawy B, Sheldon H, Arora A, Buffa FM. Single cell RNA-sequencing: A powerful yet still challenging technology to study cellular heterogeneity. Bioessays. 2022;44(11):e2200084.

18. Kogel U, Wong ET, Szostak J, Tan WT, Lucci F, Leroy P, et al. Impact of whole-body versus nose-only inhalation exposure systems on systemic, respiratory, and cardiovascular endpoints in a 2-month cigarette smoke exposure study in the ApoE(-/-) mouse model. J Appl Toxicol. 2021;41(10):1598–619.

19. Olmedo P, Goessler W, Tanda S, Grau-Perez M, Jarmul S, Aherrera A, et al. Metal Concentrations in e-Cigarette Liquid and Aerosol Samples: The Contribution of Metallic Coils. Environ Health Perspect. 2018;126(2):027010.

20. Aherrera A, Lin JJ, Chen R, Tehrani M, Schultze A, Borole A, et al. Metal Concentrations in E-Cigarette Aerosol Samples: A Comparison by Device Type and Flavor. Environ Health Perspect. 2023;131(12):127004.

21. Effah F, Shaikh SB, Chalupa D, Faizan MI, Elder A, Rahman I. High-resistance coils in E-cigarettes increase heavy metals leaching into aerosols to cause oxidants generation in human bronchial epithelial cells at air-liquid interface: A unique non-animal methodological approach on vaping studies. NAM Journal. 2025;1:100060.

22. Del Fresno C, Sancho D. Myeloid cells in sensing of tissue damage. Curr Opin Immunol. 2021;68:34–40.

23. Marshall JS, Warrington R, Watson W, Kim HL. An introduction to immunology and immunopathology. Allergy Asthma Clin Immunol. 2018;14(Suppl 2):49.

24. Lamb T, Kaur G, Rahman I. Tobacco-Derived and Tobacco-Free Nicotine cause differential inflammatory cell influx and MMP9 in mouse lung. Res Sq. 2023.

25. Wang Q, Khan NA, Muthumalage T, Lawyer GR, McDonough SR, Chuang TD, et al. Dysregulated repair and inflammatory responses by e-cigarette-derived inhaled nicotine and humectant propylene glycol in a sex-dependent manner in mouse lung. FASEB Bioadv. 2019;1(10):609–23.

26. Lamb T, Muthumalage T, Meehan-Atrash J, Rahman I. Nose-Only Exposure to Cherry– and Tobacco-Flavored E-Cigarettes Induced Lung Inflammation in Mice in a Sex-Dependent Manner. Toxics. 2022;10(8).

27. Lee PY, Wang JX, Parisini E, Dascher CC, Nigrovic PA. Ly6 family proteins in neutrophil biology. J Leukoc Biol. 2013;94(4):585–94.

28. Dannappel MV, Sooraj D, Loh JJ, Firestein R. Molecular and in vivo Functions of the CDK8 and CDK19 Kinase Modules. Front Cell Dev Biol. 2018;6:171.

29. Jin Q, Zhao J, Zhao Z, Zhang S, Sun Z, Shi Y, et al. CAMK1D Inhibits Glioma Through the PI3K/AKT/mTOR Signaling Pathway. Front Oncol. 2022;12:845036.

30. Vallin J, Grantham J. The role of the molecular chaperone CCT in protein folding and mediation of cytoskeleton-associated processes: implications for cancer cell biology. Cell Stress Chaperones. 2019;24(1):17–27.

31. Haeri M, Knox BE. Endoplasmic Reticulum Stress and Unfolded Protein Response Pathways: Potential for Treating Age-related Retinal Degeneration. J Ophthalmic Vis Res. 2012;7(1):45–59.

32. Lopes N, Galluso J, Escalière B, Carpentier S, Kerdiles YM, Vivier E. Tissue-specific transcriptional profiles and heterogeneity of natural killer cells and group 1 innate lymphoid cells. Cell Rep Med. 2022;3(11):100812.

33. Pommerenke C, Wilk E, Srivastava B, Schulze A, Novoselova N, Geffers R, et al. Global transcriptome analysis in influenza-infected mouse lungs reveals the kinetics of innate and adaptive host immune responses. PLoS One. 2012;7(7):e41169.

34. Akter S, Chauhan KS, Dunlap MD, Choreño-Parra JA, Lu L, Esaulova E, et al. Mycobacterium tuberculosis infection drives a type I IFN signature in lung lymphocytes. Cell Rep. 2022;39(12):110983.

35. Kong S, Chen TX, Jia XL, Cheng XL, Zeng ML, Liang JY, et al. Cell-specific NFIA upregulation promotes epileptogenesis by TRPV4-mediated astrocyte reactivity. J Neuroinflammation. 2023;20(1):247.

36. Hiraike Y, Saito K, Oguchi M, Wada T, Toda G, Tsutsumi S, et al. NFIA in adipocytes reciprocally regulates mitochondrial and inflammatory gene program to improve glucose homeostasis. Proc Natl Acad Sci U S A. 2023;120(31):e2308750120.

37. Sindelar JL. Regulating Vaping – Policies, Possibilities, and Perils. N Engl J Med. 2020;382(20):e54.

38. Bhalerao A, Sivandzade F, Archie SR, Cucullo L. Public Health Policies on E-Cigarettes. Curr Cardiol Rep. 2019;21(10):111.

39. Azagba S, Ebling T, Adekeye OT, Hall M, Jensen JK. Loopholes for Underage Access in E-Cigarette Delivery Sales Laws, United States, 2022. Am J Public Health. 2023;113(5):568–76.

40. Cheng D, Lee B, Jeffers AM, Stover M, Kephart L, Chadwick G, et al. State E-Cigarette Flavor Restrictions and Tobacco Product Use in Youths and Adults. JAMA Netw Open. 2025;8(7):e2524184.

41. N KEaE. Current Electronic Cigarette Use Among Adults Aged 18 and Over: United States, 2021 2023.

42. Lamb T, Muthumalage T, Rahman I. Pod-based menthol and tobacco flavored e-cigarettes cause mitochondrial dysfunction in lung epithelial cells. Toxicol Lett. 2020;333:303–11.

43. Corriden R, Moshensky A, Bojanowski CM, Meier A, Chien J, Nelson RK, et al. E-cigarette use increases susceptibility to bacterial infection by impairment of human neutrophil chemotaxis, phagocytosis, and NET formation. Am J Physiol Cell Physiol. 2020;318(1):C205–c14.

44. Jasper AE, Faniyi AA, Davis LC, Grudzinska FS, Halston R, Hazeldine J, et al. E-cigarette vapor renders neutrophils dysfunctional due to filamentous actin accumulation. J Allergy Clin Immunol. 2024;153(1):320–9.e8.

45. Ren X, Lin L, Sun Q, Li T, Sun M, Sun Z, et al. Metabolomics-based safety evaluation of acute exposure to electronic cigarettes in mice. Sci Total Environ. 2022;839:156392.

46. Kuntic M, Oelze M, Steven S, Kröller-Schön S, Stamm P, Kalinovic S, et al. Short-term e-cigarette vapour exposure causes vascular oxidative stress and dysfunction: evidence for a close connection to brain damage and a key role of the phagocytic NADPH oxidase (NOX-2). Eur Heart J. 2020;41(26):2472–83.

47. Wang Q, Sundar IK, Li D, Lucas JH, Muthumalage T, McDonough SR, et al. E-cigarette-induced pulmonary inflammation and dysregulated repair are mediated by nAChR α7 receptor: role of nAChR α7 in SARS-CoV-2 Covid-19 ACE2 receptor regulation. Respir Res. 2020;21(1):154.

48. Zhao D, Aravindakshan A, Hilpert M, Olmedo P, Rule AM, Navas-Acien A, et al. Metal/Metalloid Levels in Electronic Cigarette Liquids, Aerosols, and Human Biosamples: A Systematic Review. Environ Health Perspect. 2020;128(3):36001.

49. Zhao D, Navas-Acien A, Ilievski V, Slavkovich V, Olmedo P, Adria-Mora B, et al. Metal concentrations in electronic cigarette aerosol: Effect of open-system and closed-system devices and power settings. Environ Res. 2019;174:125–34.

50. Alcantara C, Chaparro L, Zagury GJ. Occurrence of metals in e-cigarette liquids: Influence of coils on metal leaching and exposure assessment. Heliyon. 2023;9(3):e14495.

51. Omaiye EE, Williams M, Bozhilov KN, Talbot P. Design features and elemental/metal analysis of the atomizers in pod-style electronic cigarettes. PLoS One. 2021;16(3):e0248127.

52. Rastian B, Wilbur C, Curtis DB. Transfer of Metals to the Aerosol Generated by an Electronic Cigarette: Influence of Number of Puffs and Power. Int J Environ Res Public Health. 2022;19(15).

53. Kalininskiy A, Kittel J, Nacca NE, Misra RS, Croft DP, McGraw MD. E-cigarette exposures, respiratory tract infections, and impaired innate immunity: a narrative review. Pediatr Med. 2021;4.

54. Rebuli ME, Glista-Baker E, Hoffman JR, Duffney PF, Robinette C, Speen AM, et al. Electronic-Cigarette Use Alters Nasal Mucosal Immune Response to Live-attenuated Influenza Virus. A Clinical Trial. Am J Respir Cell Mol Biol. 2021;64(1):126–37.

55. Reidel B, Radicioni G, Clapp PW, Ford AA, Abdelwahab S, Rebuli ME, et al. E-Cigarette Use Causes a Unique Innate Immune Response in the Lung, Involving Increased Neutrophilic Activation and Altered Mucin Secretion. Am J Respir Crit Care Med. 2018;197(4):492–501.

56. Song MA, Kim JY, Gorr MW, Miller RA, Karpurapu M, Nguyen J, et al. Sex-specific lung inflammation and mitochondrial damage in a model of electronic cigarette exposure in asthma. Am J Physiol Lung Cell Mol Physiol. 2023;325(5):L568–l79.

57. Chirumamilla P, Rousselle, D, Sharma, S, Commodore, S and Silveyra, P. Evaluation of Sex Differences in DNA Methylation and Lung Tissue Gene Expression in a Mouse Model of E-cigarette Exposure. Respiratory Physiology. 2024;39(S1).

58. Qiu P. Embracing the dropouts in single-cell RNA-seq analysis. Nature Communications. 2020;11(1):1169.

59. Palit S, Heuser C, de Almeida GP, Theis FJ, Zielinski CE. Meeting the Challenges of High-Dimensional Single-Cell Data Analysis in Immunology. Front Immunol. 2019;10:1515.

60. Goss K, Grant ML, Caldwell C, Dallalio GA, Stephenson ST, Fitzpatrick AM, et al. Single-cell RNA-sequencing of circulating eosinophils from asthma patients reveals an inflammatory signature. iScience. 2025;28(6):112609.

61. Jukema BN, Smit K, Hopman MTE, Bongers C, Pelgrim TC, Rijk MH, et al. Neutrophil and Eosinophil Responses Remain Abnormal for Several Months in Primary Care Patients With COVID-19 Disease. Front Allergy. 2022;3:942699.

62. Flinkman E, Vähätalo I, Tuomisto LE, Lehtimäki L, Nieminen P, Niemelä O, et al. Association Between Blood Eosinophils and Neutrophils With Clinical Features in Adult-Onset Asthma. The Journal of Allergy and Clinical Immunology: In Practice. 2023;11(3):811–21.e5.

63. Lourda M, Dzidic M, Hertwig L, Bergsten H, Palma Medina LM, Sinha I, et al. High-dimensional profiling reveals phenotypic heterogeneity and disease-specific alterations of granulocytes in COVID-19. Proceedings of the National Academy of Sciences. 2021;118(40):e2109123118.

64. Deniset JF, Surewaard BG, Lee WY, Kubes P. Splenic Ly6G(high) mature and Ly6G(int) immature neutrophils contribute to eradication of S. pneumoniae. J Exp Med. 2017;214(5):1333–50.

65. Kleinholz CL, Riek-Burchardt M, Seiß EA, Amore J, Gintschel P, Philipsen L, et al. Ly6G deficiency alters the dynamics of neutrophil recruitment and pathogen capture during Leishmania major skin infection. Scientific Reports. 2021;11(1):15071.

66. Sprenkeler EGG, Zandstra J, van Kleef ND, Goetschalckx I, Verstegen B, Aarts CEM, et al. S100A8/A9 Is a Marker for the Release of Neutrophil Extracellular Traps and Induces Neutrophil Activation. Cells. 2022;11(2).

67. Guo Q, Zhao Y, Li J, Liu J, Yang X, Guo X, et al. Induction of alarmin S100A8/A9 mediates activation of aberrant neutrophils in the pathogenesis of COVID-19. Cell Host & Microbe. 2021;29(2):222–35.e4.

68. Averill MM, Kerkhoff C, Bornfeldt KE. S100A8 and S100A9 in cardiovascular biology and disease. Arterioscler Thromb Vasc Biol. 2012;32(2):223–9.

69. Wang S, Song R, Wang Z, Jing Z, Wang S, Ma J. S100A8/A9 in Inflammation. Front Immunol. 2018;9:1298.

70. Mair I, Wolfenden A, Lowe AE, Bennett A, Muir A, Smith H, et al. A lesson from the wild: The natural state of eosinophils is Ly6G(hi). Immunology. 2021;164(4):766–76.

71. Shanker A, Marincola FM. Cooperativity of adaptive and innate immunity: implications for cancer therapy. Cancer Immunol Immunother. 2011;60(8):1061–74.

72. Carroll MC, Prodeus AP. Linkages of innate and adaptive immunity. Curr Opin Immunol. 1998;10(1):36–40.

73. Wang R, Zhang H, Xu J, Zhang N, Pan T, Zhong X, et al. MALT1 Inhibition as a Therapeutic Strategy in T-Cell Acute Lymphoblastic Leukemia by Blocking Notch1-Induced NF-κB Activation. Front Oncol. 2020;10:558339.

74. Bird CH, Christensen ME, Mangan MS, Prakash MD, Sedelies KA, Smyth MJ, et al. The granzyme B-Serpinb9 axis controls the fate of lymphocytes after lysosomal stress. Cell Death Differ. 2014;21(6):876–87.

75. Beland M, Desjardins M, Xue D, Mazer BD. Semaphorin 4C Is An Intrinsic Regulator Of Cell-Cell Interaction In Th2 Stimulated Memory-B-Cells. Journal of Allergy and Clinical Immunology. 2014;133(2):AB249.

76. Szilagyi Z, Gustafsson CM. Emerging roles of Cdk8 in cell cycle control. Biochimica et Biophysica Acta (BBA) – Gene Regulatory Mechanisms. 2013;1829(9):916–20.

77. Brackley KI, Grantham J. Activities of the chaperonin containing TCP-1 (CCT): implications for cell cycle progression and cytoskeletal organisation. Cell Stress Chaperones. 2009;14(1):23–31.

78. Berry R, Ng N, Saunders PM, Vivian JP, Lin J, Deuss FA, et al. Targeting of a natural killer cell receptor family by a viral immunoevasin. Nat Immunol. 2013;14(7):699–705.

79. Bolanos FD, Tripathy SK. Activation receptor-induced tolerance of mature NK cells in vivo requires signaling through the receptor and is reversible. J Immunol. 2011;186(5):2765–71.

80. Israr M, DeVoti JA, Papayannakos CJ, Bonagura VR. Role of chemokines in HPV-induced cancers. Seminars in Cancer Biology. 2022;87:170–83.

81. Li BH, Garstka MA, Li ZF. Chemokines and their receptors promoting the recruitment of myeloid-derived suppressor cells into the tumor. Mol Immunol. 2020;117:201–15.

82. Wang S, Sun Y, Li C, Chong Y, Ai M, Wang Y, et al. TH1L involvement in colorectal cancer pathogenesis by regulation of CCL20 through the NF-κB signalling pathway. J Cell Mol Med. 2024;28(10):e18391.

83. Ross SH, Cantrell DA. Signaling and Function of Interleukin-2 in T Lymphocytes. Annu Rev Immunol. 2018;36:411–33.

84. Rapp M, Wintergerst MWM, Kunz WG, Vetter VK, Knott MML, Lisowski D, et al. CCL22 controls immunity by promoting regulatory T cell communication with dendritic cells in lymph nodes. J Exp Med. 2019;216(5):1170–81.

85. Jinquan T, Quan S, Feili G, Larsen CG, Thestrup-Pedersen K. Eotaxin activates T cells to chemotaxis and adhesion only if induced to express CCR3 by IL-2 together with IL-4. J Immunol. 1999;162(7):4285–92.

86. Chia TY, Billingham LK, Boland L, Katz JL, Arrieta VA, Shireman J, et al. The CXCL16-CXCR6 axis in glioblastoma modulates T-cell activity in a spatiotemporal context. Front Immunol. 2023;14:1331287.

87. Böttcher JP, Beyer M, Meissner F, Abdullah Z, Sander J, Höchst B, et al. Functional classification of memory CD8+ T cells by CX3CR1 expression. Nature Communications. 2015;6(1):8306.

88. Tu GW, Ju MJ, Zheng YJ, Hao GW, Ma GG, Hou JY, et al. CXCL16/CXCR6 is involved in LPS-induced acute lung injury via P38 signalling. J Cell Mol Med. 2019;23(8):5380–9.

89. Christiani DC. Vaping-Induced Acute Lung Injury. N Engl J Med. 2020;382(10):960–2.

90. Kagohashi K, Sasaki Y, Ozawa K, Tsuchiya T, Kawahara S, Saitoh K, et al. Role of Signal-Transducing Adaptor Protein-1 for T Cell Activation and Pathogenesis of Autoimmune Demyelination and Airway Inflammation. The Journal of Immunology. 2024;212(6):951–61.

91. Zhu X, Guan R, Zou Y, Li M, Chen J, Zhang J, et al. Cold-inducible RNA binding protein alleviates iron overload-induced neural ferroptosis under perinatal hypoxia insult. Cell Death & Differentiation. 2024;31(4):524–39.

92. Firsanov D, Zacher M, Tian X, Sformo TL, Zhao Y, Tombline G, et al. Evidence for improved DNA repair in long-lived bowhead whale. Nature. 2025.

93. Mikulski Z, Zaslona Z, Cakarova L, Hartmann P, Wilhelm J, Tecott LH, et al. Serotonin activates murine alveolar macrophages through 5-HT2C receptors. Am J Physiol Lung Cell Mol Physiol. 2010;299(2):L272–80.

94. Dutta S, Sengupta P. Men and mice: Relating their ages. Life Sci. 2016;152:244–8.

95. Jackson SJ, Andrews N, Ball D, Bellantuono I, Gray J, Hachoumi L, et al. Does age matter? The impact of rodent age on study outcomes. Lab Anim. 2017;51(2):160–9.

96. Lee YO, Nonnemaker JM, Bradfield B, Hensel EC, Robinson RJ. Examining Daily Electronic Cigarette Puff Topography Among Established and Nonestablished Cigarette Smokers in their Natural Environment. Nicotine Tob Res. 2018;20(10):1283–8.

97. Kaur GR, I. URMC TriState SenNet Mouse Lung Digestion. protocols.io 2024.

98. Hao Y, Hao S, Andersen-Nissen E, Mauck WM, 3rd, Zheng S, Butler A, et al. Integrated analysis of multimodal single-cell data. Cell. 2021;184(13):3573–87.e29.

99. Schaum N, Karkanias J, Neff NF, May AP, Quake SR, Wyss-Coray T, et al. Single-cell transcriptomics of 20 mouse organs creates a Tabula Muris. Nature. 2018;562(7727):367–72.

100. Hurskainen M, Mižíková I, Cook DP, Andersson N, Cyr-Depauw C, Lesage F, et al. Single cell transcriptomic analysis of murine lung development on hyperoxia-induced damage. Nat Commun. 2021;12(1):1565.

101. Yu G, Wang LG, Han Y, He QY. clusterProfiler: an R package for comparing biological themes among gene clusters. Omics. 2012;16(5):284–7.

