## Supplementary material for "Single-cell transcriptomics identifies altered neutrophil dynamics and accentuated T-cell cytotoxicity in tobacco flavored e-cigarette exposed mouse lungs": suppl text and figs

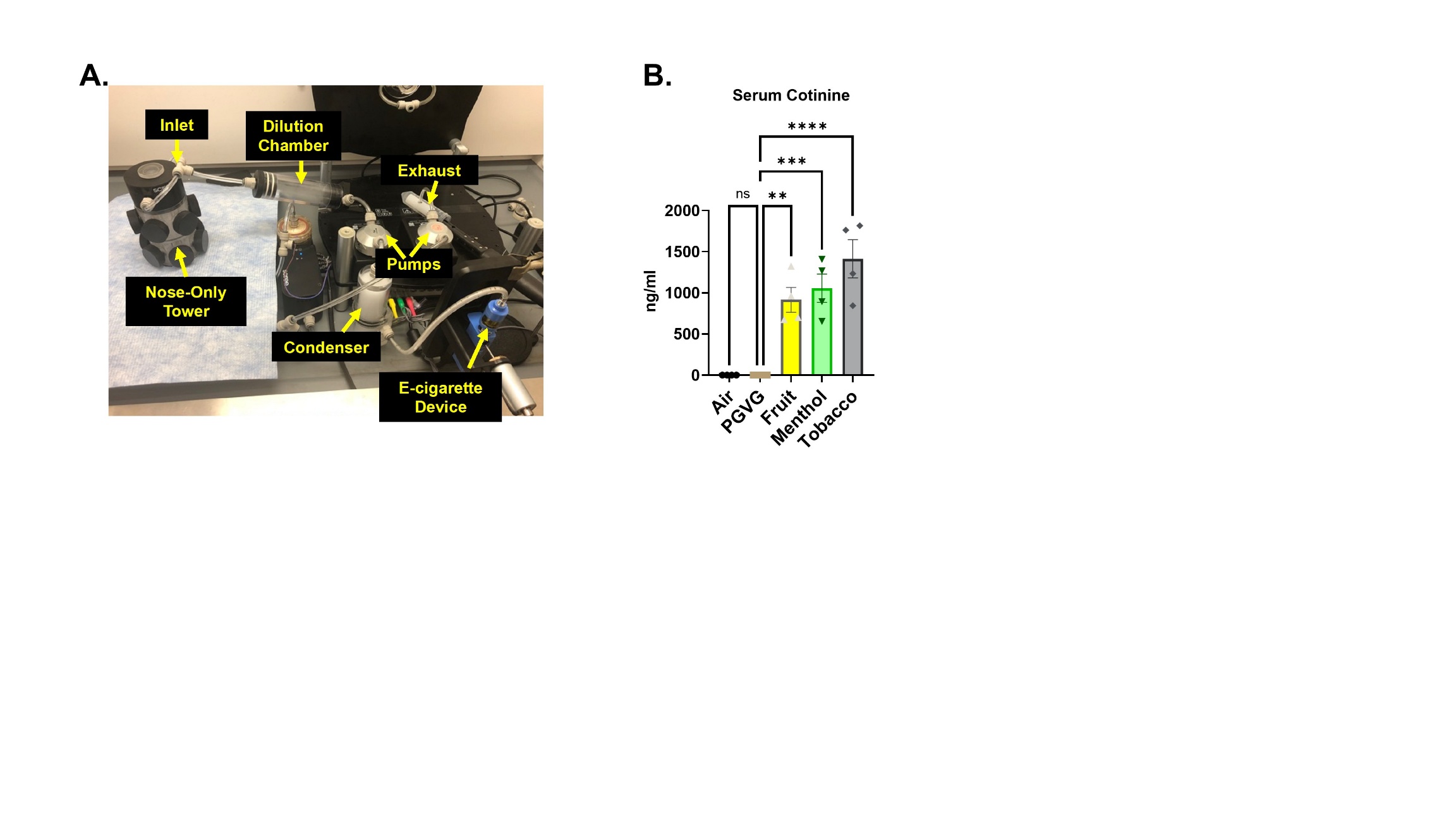

**Figure S1:** The nose only exposure system used for performing the mouse experiment **(A)**. The exposure characteristics were assessed by measuring the serum cotinine levels in the blood of exposed and control mice. Data are shown as mean ± SEM (n = 4/group); ns: not significant. **p<0.01, ***p<0.001 and **** p< 0.0001 vs air, per one-way ANOVA for multiple comparison **(B)**.

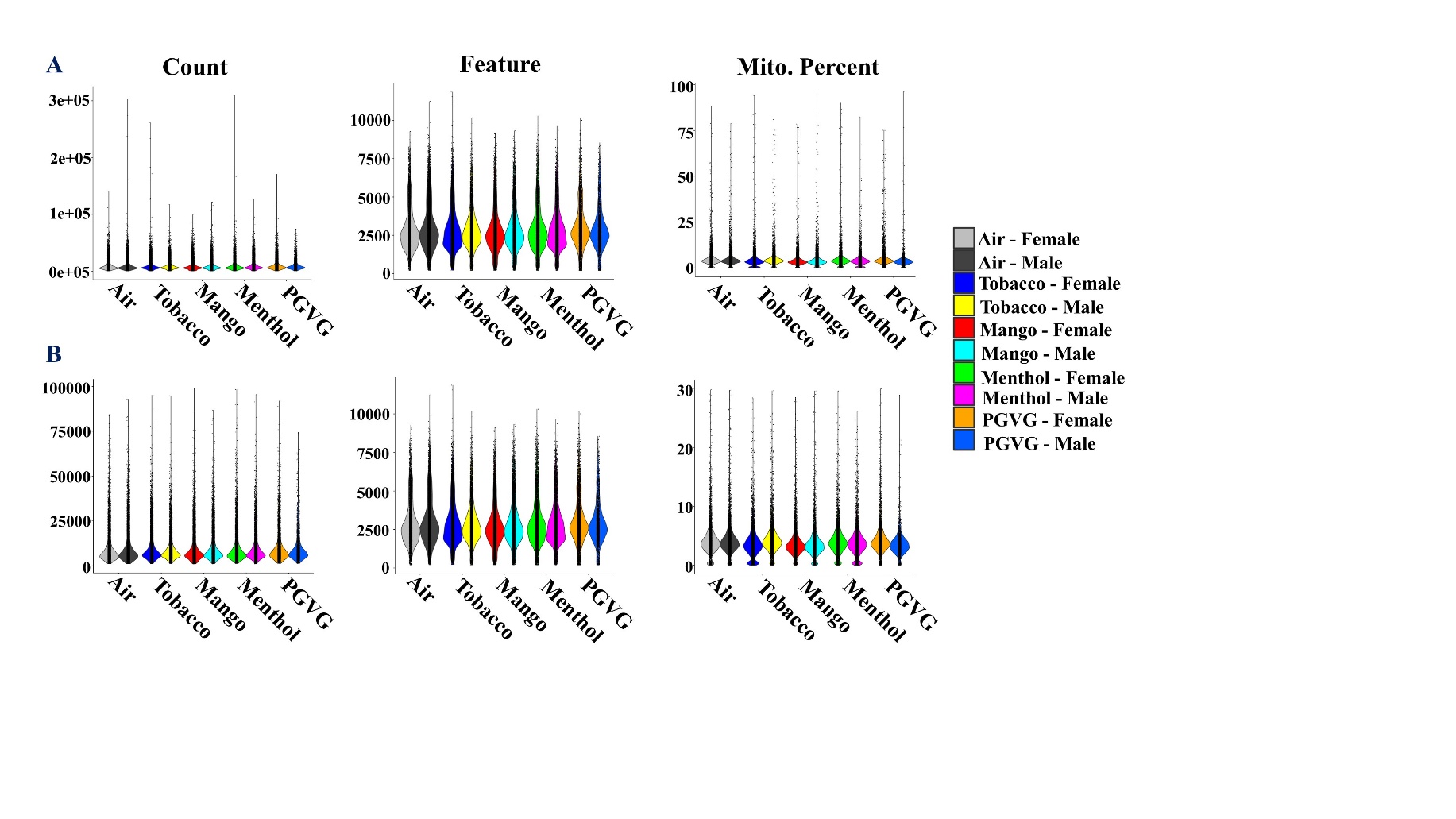

**Figure S2:** Figure showing the normalized counts, features and mitochondrial gene percentage in the integrated single call data before **(A)** and after **(B)** normalization.

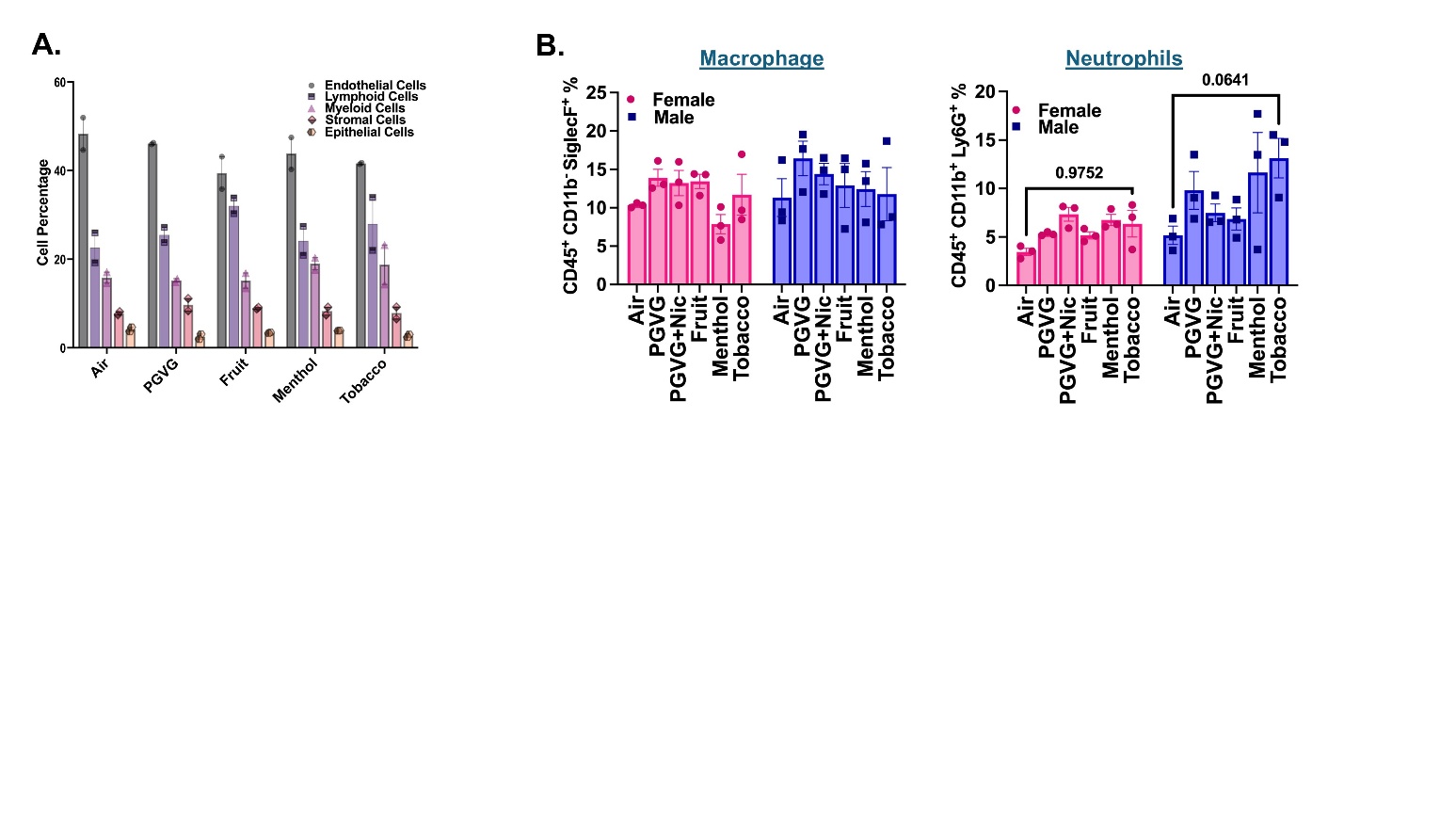

**Figure S3:** Cell frequencies of major cell clusters (epithelial, endothelial, stromal, myeloid and lymphoid) in control (air) and exposed (PGVG, fruit, menthol and tobacco) as determined by scRNA seq **(A).** Flow cytometry results showing changes in relative percentages of macrophages and neutrophils **(B)** in the lung digests from exposed (PGVG, PGVG + Nic, Fruit, Menthol and tobacco) and control (Air) mice following 5-day acute exposure. The data represented in this figure is an extension of the data represented in Figure 3C-D of the main manuscript with the addition of an extra group (PGVG + Nic).

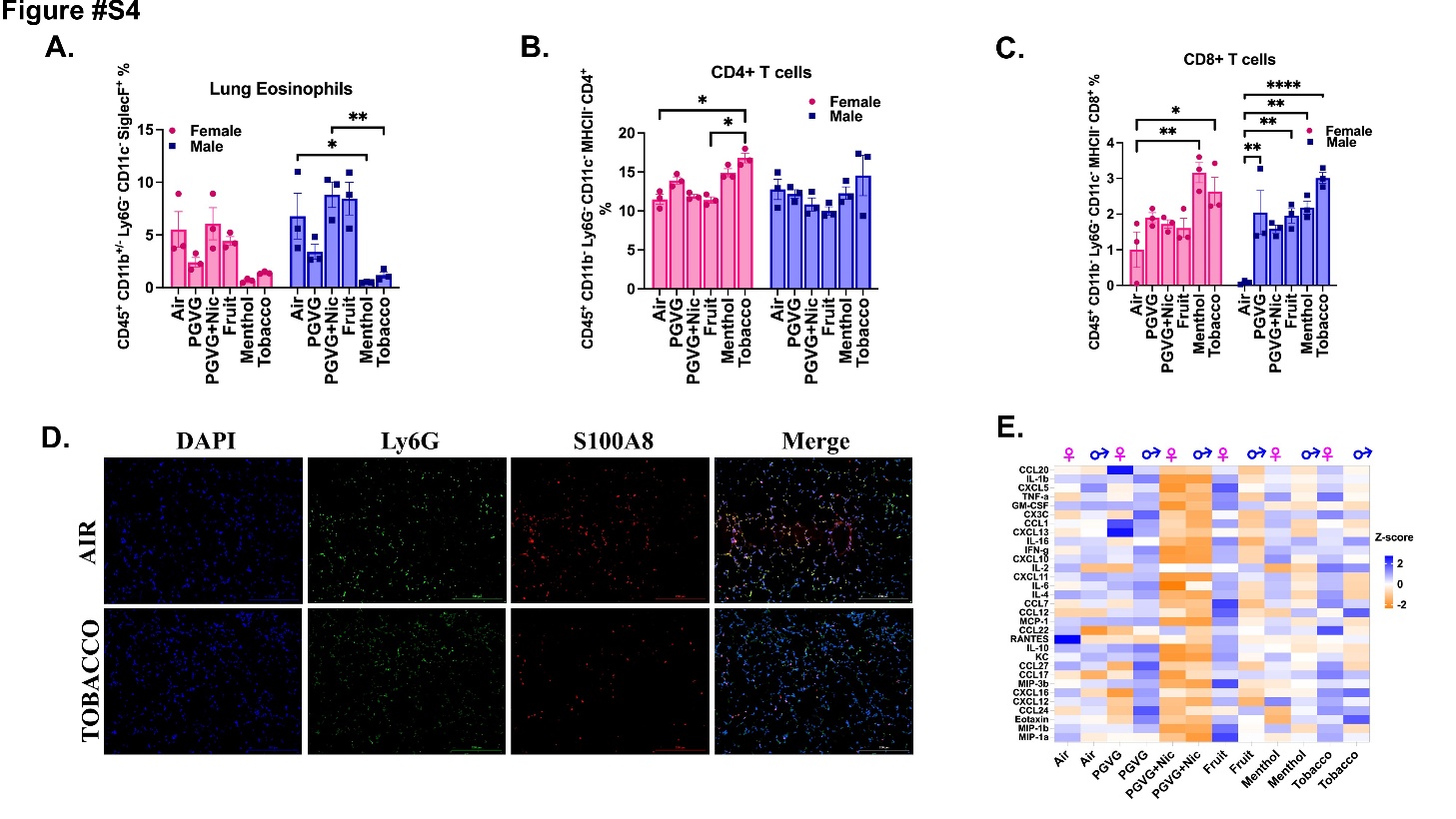

**Figure S4:** Flow cytometry results showing the changes in the mean counts of eosinophils **(A)** CD4+ **(B)** and CD8+ **(C)** T-cells in the lung digests from exposed (PGVG, PGVG + Nic, Fruit, Menthol and tobacco) and control (Air) mice following 5-day acute exposure. The data represented in this figure is an extension of the data represented in Figure 4 and 5C-D of the main manuscript with the addition of an extra group (PGVG + Nic). Co-immunofluorescence results showing single channel staining for DAPI (blue channel), Ly6G (green channel) and S100A8 (red channel) along with the merges images in lung tissue sections from control and tobacco-flavored e-cig aerosol treated mouse lungs. The above image is an extension of the data presented in Figure 6C-D of the main manuscript **(D)**. The levels of pro-inflammatory cytokines/chemokines in the lung digests from experimental (PGVG, PGVG+Nic, Fruit, Menthol and Tobacco) and control (air) were assessed using multianalyte assay. The results obtained were plotted as a heatmap with the z-scores represented between the scale of orange (low) to blue (high) **(E)**.

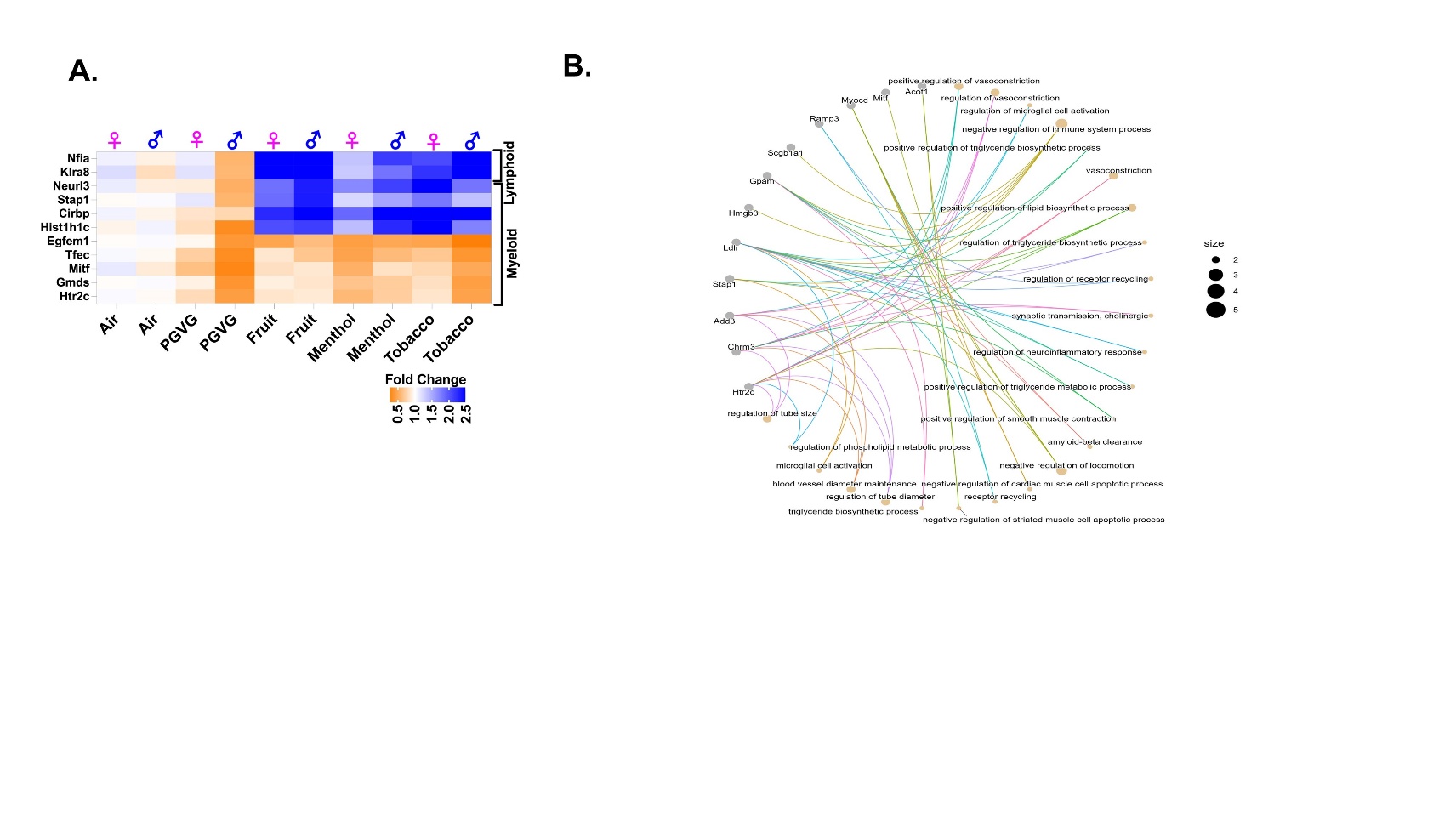
**Figure S5:** Heatmap showing the fold changes in the expression of commonly dysregulated genes in the myeloid and lymphoid clusters in mouse lungs exposed to flavored e-cig aerosols as compared to ambient air as determined after DESeq2 analyses **(A)**. CNET plot results showing the pathways regulated by the common DEGs on acute (5-day) exposure to differently flavored (fruit, menthol, and tobacco) e-cig aerosols in C57BL/6J mouse lungs **(B)**.

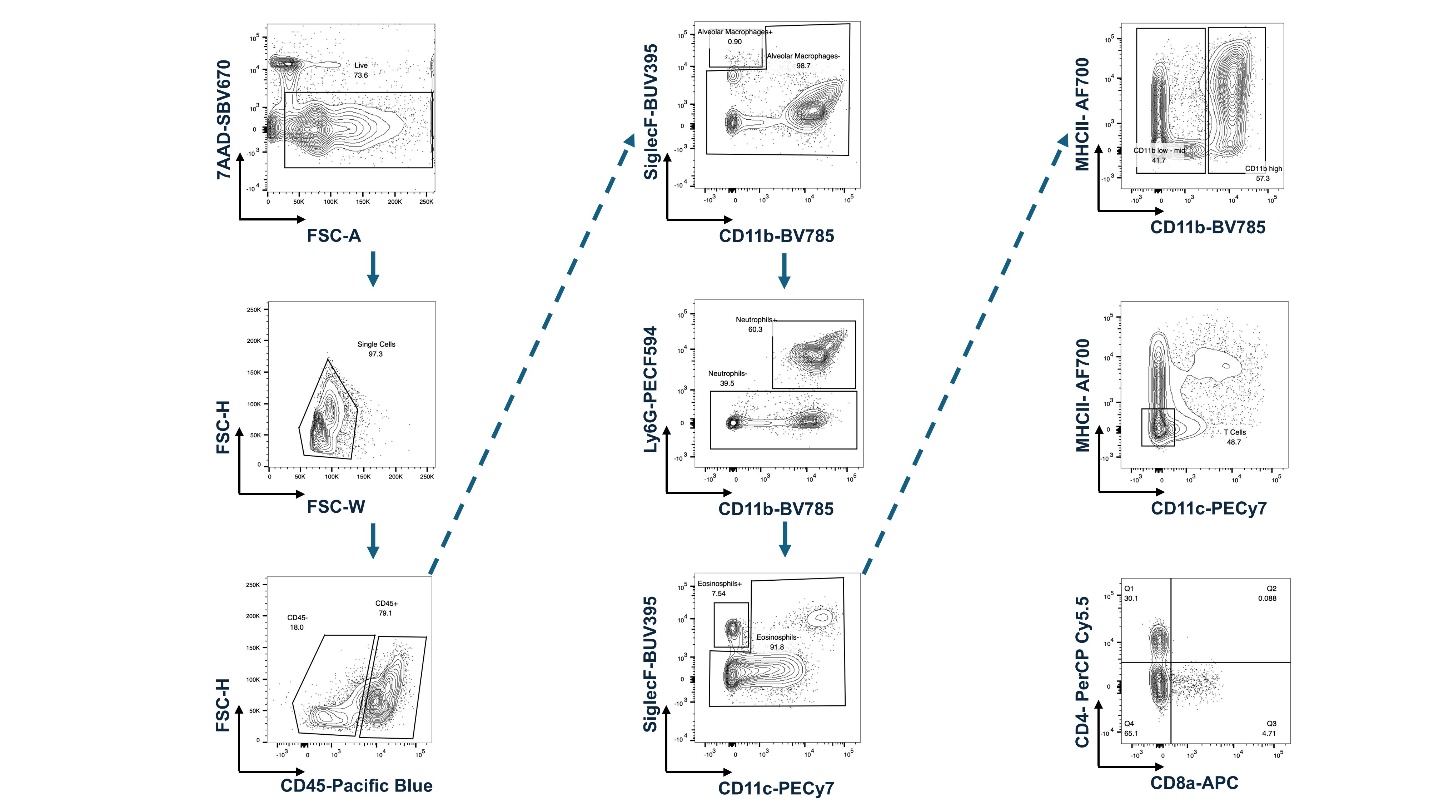

**Figure S6:** Representative plots showing the gating strategy used to gate for macrophages, neutrophils, eosinophils, CD4+ and CD8+ T cells using flow cytometry. Data is analyzed using FlowJo software.
